# Xtricorder: A likelihood-enhanced self-rotation function and application to a machine-learning enhanced Matthews prediction of asymmetric unit copy number

**DOI:** 10.1101/2025.05.22.655506

**Authors:** Airlie J. McCoy, Randy J. Read

## Abstract

Analysis of crystallographic diffraction data before phasing gives the crystallographer a ‘first look’ at the nature of the problem and the context in which the structure determination will be performed. We here report the development of *Xtricorder*, an application that targets analysis of crystallographic data specifically for likelihood-based phasing. As well as porting many of the analyses previously available but relatively inaccessible in our *Phaser* codebase, *Xtricorder* offers a likelihood-enhanced self-rotation function. A novel and intuitive graphical representation of the self-rotation function presents the results for user inspection, and has the added advantage that, in an adapted form, is appropriate for training a convolutional neural network to enhance the standard Matthews analysis and more accurately predict the number of copies in the asymmetric unit. We investigate the usefulness of the likelihood-enhanced self-rotation function in ‘first look’ analyses, exploring the circumstances under which the self-rotation function results are useful, and discuss the application to AI-generated structure prediction.

**Synopsis** *Xtricorder* is a new tool for analysing crystallographic data prior to phasing, featuring a likelihood-enhanced self-rotation function and graphical output that aids both user interpretation and machine learning-based prediction of asymmetric unit content.

## 1. Introduction

*Xtricorder* for crystallographic data analysis builds on software already available that has been developed both by our group and others. Indeed, the name is partly inspired by the name of *Xtriage* (Zwart *et al*., 2005), a component of the *PHENIX* suite (Liebschner *et al*., 2019), which performs diagnostic tests for crystallographic structure solution. *Xtriage* assesses the overall completeness, redundancy, and the presence of possible pathologies like anisotropy, translational non-crystallographic symmetry (tNCS), twinning and pseudo-symmetries and has been extensively cited as a useful ‘first look’ tool.

The *CCP4* suite (Agirre *et al*., 2023; Winn *et al*., 2011) has programs that fulfil a similar role, although not wrapped in a single application: *Ctruncate* for anisotropy analysis and twinning tests (French & Wilson, 1978), *Pointless* for space group analysis (Evans, 2006), *Zanuda* for space group subgroup analysis (Lebedev & Isupov, 2014), and *Auspex* for visual inspection of data intensity pathologies (Thorn *et al*., 2017). In addition, *CCP4* has two implementations of a self-rotation function, one in *Molrep* (Vagin & Teplyakov, 2010) and one in *Polarrfn* (Agirre *et al*., 2023; Winn *et al*., 2011).

Our software *Phaser* (McCoy *et al*., 2007) , distributed with both *Phenix* and *CCP4*, performs an array of analyses that are necessary/optimal for application of the maximum likelihood functions for phasing by molecular replacement (MR), single-wavelength anomalous dispersion (SAD) and the combined MR-SAD method. These also offer useful ‘first look’ analyses: correction of systematic intensity modulations due to anisotropy and translational non-crystallographic symmetry (tNCS) (Caballero *et al*., 2020; Read *et al*., 2013); cell content analysis to estimate the fraction scattering of atoms/models for the σ_A_ estimation (Read, 1986); and twinning tests (Padilla & Yeates, 2003) particularly in the presence of tNCS. The presence of twinning modifies the target for the expected-log-likelihood gain directed optimization of the resolution for phasing (Oeffner *et al*., 2018). Additionally, reflections are analysed for information content and the set of reflections is filtered to remove those reflections that only slow down the calculations.

The reorganisation, modifications and expansion of the codebase on moving from *Phaser* to *Phasertng* (McCoy *et al*., 2020) offered the opportunity to not only make this functionality available in a dedicated stand-alone application but also offered the opportunity to add a self-rotation function (SRF). By wrapping these functionalities, we aim to make them more accessible and visible to crystallographers, so as to encourage uptake.

The SRF provides insight into rotational relationships within the crystal lattice including non-crystallographic components. Historically, it has been used on an *ad hoc* basis to discern the point group symmetry of oligomers and indications of the number of copies in the asymmetric unit (Rossmann & Blow, 1962). However, the SRF is not currently standardly applied in structure solution pipelines, nor automatically mined for information pertinent to structure solution *a priori*.

Any internal symmetry of an oligomer is known to be an advantage during crystallization (Chruszcz *et al*., 2008) and has even been proposed as a mechanism to improve crystallizability (Banatao *et al*., 2006). Many proteins form oligomers in solution with a point group symmetry that is capable of being represented by a crystal symmetry (e.g. C_3_ trimers in P3, D_2_ tetramers in P222), however during crystallization these point groups are not always incorporated as crystallographic symmetry. Likewise, an oligomeric point group symmetry for which a *subgroup* of the point group may be represented by crystal symmetry (e.g. C_12_ in P6), may or may not be *partly* crystallographic. Conversely, proteins that are not oligomeric in solution may crystallize with non-crystallographic symmetry (as well as crystallographic symmetry). Therefore, even when the SRF gives a clear signal for symmetry, the relationship between the SRF identified symmetry and the oligomeric associations and asymmetric unit composition is far from obvious or straightforward. Biophysical analysis of oligomerization state is invaluable in this regard.

In the era of AI-generated structure prediction, the SRF also has the potential to provide useful information about oligomeric associations for structure prediction. In the absence of a structure, determination of the oligomeric state can direct structure prediction protocols to target a particular oligomeric state and thereby improve predictions, which may tip the scales in improving the model for molecular replacement phasing. It is therefore of interest to look again at the information that can be provided by the SRF in a systematic study.

## 2. Self-Rotation Function

The SRF identifies rotational relationships in the crystal, including symmetries, by comparison of the data against itself. Only point group symmetry is detected in this analysis: any translations associated with the rotations are not identified (for example 2_1_ screw rotations are reduced to 2-fold rotations). Crystallographic and non-crystallographic point groups are described in Section 10.1 of the International Tables for Crystallography (Hahn & Klapper, 2006).

SRFs have long been implemented in formerly popular programs *X-PLOR* (Brünger, 1992) and *CNS* (Brunger, 2007), and currently popular programs *Molrep* and *Polarrfn*. These implementations use the Patterson function, which is the Fourier transform of the intensities, and which gives a vector map of the crystal (for review see (Evans and McCoy, 2008)). Vectors close to the origin represent intra-molecular associations and associations at crystal contacts, while at increasing distances from the origin, increasing proportions of vectors represent inter-molecular relationships. Cutting out a sphere around the Patterson origin enriches the Patterson for intra-molecular vectors.

Since rotations of molecules are echoed in the rotations of the corresponding vectors, the molecular rotations are identified by a match of the radius-restricted Patterson against a copy of itself after rotation. The match can be measured as various functions, such as a product function or a correlation coefficient, or as the equivalent to the Patterson product function in reciprocal space. The reciprocal-space version can be made fast by factorization and FFT and is known as the ‘fast rotation function’ (Navaza, 1994). The highest peak is at 0° rotation and the height of other peaks is normally measured as the percentage of this (100%) ‘origin’ peak, with the mean taken as 0%. In this way, the SRF is analysed analogously to the way the native Patterson is used to identify tNCS, where non-origin peaks representing molecular translations are identified by their height relative to the ‘origin’ peak (100%) which corresponds with the molecule mapped to itself (Caballero *et al*., 2020).

## 3. Likelihood-enhanced Self Rotation Function

The established fast rotation function in *Phaser* is based on the likelihood-enhanced rotation function (LERF, (Storoni *et al*., 2004)). A first and second order approximation of the full likelihood rotation function were developed as LERF1 and LERF2. LERF1 was found to be faster (requiring only one FFT rather than two) and sufficient to find the correct orientation, and LERF2 was consigned to developer use only. Only LERF1 has been ported to *Phasertng*.

In the context of the rotation function of model against data used for molecular replacement (the cross-rotation function, CRF), LERF1 has the major advantage over the Patterson rotation function of being able to incorporate information from a fixed partial model and hence substantially enhance the signal for second and subsequent asymmetric unit model components when the asymmetric unit is built up by addition, a hallmark of the maximum likelihood approach. Where there is no fixed model and as originally published, LERF1 for the CRF reduces to the fast rotation function proposed by Bricogne (Bricogne, 1997). The coefficients for the data component are *E*_O2_ – 1, where *E*_O_ is the normalized observed structure factor amplitude, while the coefficients for the model component are σ_A_ ^2^(*E*_C2_ – 1), where *E*_C_ is the normalized calculated structure factor amplitude (from the model) and σ_A_ is the estimated correlation between the true and calculated structure factor amplitudes. These coefficients correspond to sharpened, variance weighted, origin-removed Pattersons.

Since the publication of LERF, we have developed the Log-Likelihood Gain on Intensity (LLGI) target for molecular replacement which has improved the coefficients for LERF1; LLGI-LERF1 has *E*_O_ replaced by *D_obs_E*_eff_ (Read & McCoy, 2016). These coefficients incorporate the error in the observed intensity into the likelihood target through a transformation that preserves the correct normalization of the data, particularly for data with low I/σI, avoiding the distortions introduced using the ‘inflation of the variance’ method (for review see (McCoy, 2004)). For the likelihood-enhanced SRF (LESRF), the coefficients are *D_obs_*^2^(*E*_eff_ ^2^ – 1). The LESRF is highly analogous to the LERF1 (CRF) and no further computational tools are required.

An advantage of using the LESRF over the Patterson-based SRF is a reduced dependency on the resolution of the data because data with low I/σI are intrinsically down weighted through low values of *D_obs_*. The LESRF also includes the anisotropy correction and tNCS correction terms in the calculation of *E_eff_*.

## 4. Graphical Representations

The fast SRF (like the fast CRF) results in a three-dimensional map calculated in Euler angle space, with peaks at the positions of both the crystallographic and non-crystallographic rotations. *Xtricorder* follows the tradition of other SRF software by projecting the contours for visualization using polar (stereographic) plots. The Euler angles of the calculation space are refactored as axis (φ and ψ) and angle rotations (κ), with a polar plot for each angular κ-section. The contour plot for each κ-section (range 0°-180°, normally at intervals of between 1° and 5°) allows the user to identify peaks that represent the direction of the axis of the rotation, if any. In *Xtricorder*, the SRF has a minimal sampling of 3° independent of resolution or molecular radius (unlike the implementation of the CRF), to always to sample κ-sections associated with C_n_ rotations, such as two-fold, three-folds, and four-folds, effectively.

### 4.1. Scatter Plot

The SRF in *Xtricorder* is reported as a novel scatter plot. Marker colour, shape and size are used to indicate SRF peak properties, as described below. This single plot facilitates the identification of significant rotations and their directions and reveals the patterns of crystallographic and non-crystallographic symmetry operations.

Marker size is used to indicate the peak height relative to the original peaks, where the top peaks is 100% and the mean is 0%. Peak height corresponds to marker area. Crystallographic symmetry operations will generate peak heights ∼100% of the origin peak, as will very strong non-crystallographic symmetries. The default minimal peak height selected for plotting is peaks over 5%.

To reduce noise, a further selection is applied scaling the peak heights so that the top non-crystallographic peak is taken as 100^%^, where the superscript is used to distinguish this score from the global 100% maximum. Note that the top global maximum peak height of 100% will be greater than or equal to 100^%^, possibly much greater, if no strong non-crystallographic symmetry is present. Peaks below 30^%^ are not displayed. For cases with strong non-crystallographic symmetry, this means that the effective minimal peak height is 30% not 5%.

Marker colour is used to distinguish the nearest rotational order 360°/*n* for *n* in range 2 to 12: C_2_ (red), C_3_ (gold), C_4_ (orange), C_5_ (light green), C_6_ (forest green), C_7_ (cyan), C_8_ (blue), C_9_ (plum), C_10_ (magenta), C_11_ (burlywood) and C_12_ (brown). Rotational order 360°/*n* for *n* ≥ 13 are marked with black and discussed further below (colours as defined in matplotlib (Hunter, 2007)).

Marker shape is used to distinguish between the ‘exactness’ of the rotational symmetry. Crystallographic symmetry is marked with stars. Non-crystallographic rotational symmetries 360°/*n* for *n* in range 2-12 are marked with circles if *n* is an integer (with a small tolerance) and a triangle if not.

Higher order rotations 360°/*n* for *n* ≥ 13 are uncommon. The ability to distinguish the precise order *n* of the rotation becomes increasingly difficult as *n* increases; real variation in the orientation of monomers within higher order multimers in the crystal can degrade the signal. These rotations are also less likely to correspond with C_n_ symmetry. Consequently, these rotations are all are displayed with a black marker without further distinction of *n* by colour. Likewise, the shape of the marker does not distinguish how close *n* is to an integer, unlike the cases for *n* in range 2-12. Rather, the shape is repurposed to display the value of *n* for *n* in range 13-16: inverted triangles (*n =* 13), squares (*n =* 14), pentagons (*n =* 15), and hexagons (*n =* 16). Rotations 360°/*n* for *n* in range 17-23 are displayed as black crosses, with details in the logfile’s table output. Peaks corresponding to proper rotations 360°/*n* for *n >* 24 (rotations of < 15°) are not displayed but can be found in the logfile’s table output.

The scatter plots are displayed as stereographic and Mercator plots. An example plot for a 2-ring catenane comprising two interlocking dodecameric toroids (1zye, Figure S1) is shown in Figure S2.

### 4.2. Contour Plots

The SRF in *Xtricorder* is also rendered as traditional contour plots, offering a complementary perspective that emphasizes the sharpness/shallowness of the SRF peaks selected for the scatter plot. The scatter plot representation described above is superimposed on these contours, with peaks corresponding to different ClZ rotational symmetries separated onto their corresponding κ-section. This view, with the scatter plot points distributed to different layers, gives clarity where there are overlapping points in the single scatter plot representation, and ensures that the plots remain interpretable to crystallographers with colour vision deficiencies.

Cross-referencing the contour plots with the overlaid scatter plots enables more nuanced interpretation of peak features, allowing crystallographers to distinguish between genuinely discrete rotational peaks and shoulders or extensions of broader features.

The contour plots for selected layers of 1zye are shown in Figure S3.

### 4.3. Machine Learning Representations

The graphical stereographic plot designed for user interpretation of the SRF results was adapted for machine learning, by removing constant elements of the plot (the axes and axes labels and annotations), using greyscale, and outputting images of size 112×112 pixels. This size is one quarter the widely adopted size in machine learning for image classification of 224×224 pixels, as the features of the SRF plots are sparse. This size balances preserving sufficient detail while keeping the final weights file small.

Rather than using colour for rotational order on one plot, the rotational symmetries C_2_-C_12_ were output on 11 different images, to separate markers that overlap on the combined scatter plot. Circles and triangles indicated the relationship of the rotation to circular symmetry C_2_-C_12_ as above, however the low resolution meant that greyscale was also used to distinguish the two – circles in black and triangles in grey. Size indicated peak height. Rotational symmetry over C_13_ was output on image 12 with inverted triangles, square, pentagons, hexagons and crosses as above.

In addition to the 12 SRF layers, an additional image encoded the Matthews coefficient probabilities and space group. The probabilities were output in the form of a greyscale heat map, with the intensity of the marker at each position indicating the probability. The space group was encoded as a shape with numbers for the space group point group superimposed. Information about the most prominent C_n_ symmetry, and any D_n_, O_t_ and I_h_ symmetries, were also encoded on this image.

The machine learning images for 1zye are shown in Figure S4.

## 5. Dataset generation

To test the properties of the LESRF, a dataset of structures with and without multiple copies in the asymmetric unit was generated from the PDB archive and probed to understand the dependency of the SRF on the radius of integration and the relationship between SRF-identified rotational symmetries and asymmetric unit contents.

### 5.1. Sampling of asymmetric unit copy number and space groups

A search of the PDB was performed to identify structures with up to 24 protein monomers or heterodimers (referred to henceforth as the ‘assembly’) in the asymmetric unit. The number of cases counted decreased rapidly with increasing numbers in the asymmetric unit, and even numbers were more prevalent than odd numbers. Previous analysis of the PDB revealed the same trends (Berman *et al*., 2013). A separate search was performed for the set of space groups in each of the 11 point groups, to evenly sample space group point groups (SGPG). A limit of 100 test cases per SGPG prevented vast overrepresentation of some asymmetric unit/space group point group combinations.

Additional constraints on the selection of test cases were that they had data deposited and R-factors reproducible with *PDB_REDO* (Joosten *et al*., 2012), and with a resolution of diffraction of at least 3 Å. Likewise, the search was performed separately for proteins under 100 amino acids and over 100 amino acids, to ensure a range of molecular sizes. Structures with DNA and RNA were excluded as these have strong internal rotational symmetries. Structures consisting of only coiled coils with a length to breadth ratio over 8 were excluded also because of the strong internal two-fold symmetry. Virus (capsids) were excluded as these are special cases, often having extremely low solvent content, and often having icosahedral symmetry, which is highly unlikely to be unknown to the crystallographer before crystallographic investigation. The search returned representative structures with a sequence similarity cutoff of 90%.

Data and structures for each identified PDB entry were downloaded from *PDB_REDO*, and the coordinates of one assembly (for example, chain A in the case of monomers) were extracted to a separate PDB file. Various annotations associated with the PDB entry were extracted to a database: from the PDB archive came the title, space group, resolution of the diffraction data, number of chains, number of protein entities, and number of residues; from *PDB_REDO* came the presence of twinning (as recorded in PDB header after *PDB_REDO* refinement); and from *Xtricorder* came the solvent content for the deposited structure, data anisotropy measured as ΔB in intensities between strongest and weakest diffraction directions and the model sphericity measured as the ratio of the largest to shortest extent of the assembly.

The number of cases for seven, nine and eleven in the asymmetric unit were supplemented for the machine learning studies by applying allowed reindexing operations to the data, which had the effect of changing the representation of the SRF peak positions (there were only minor computational differences). In addition, five structures with seven in the asymmetric unit for which experimental data was not available were included by using calculated data.

Histograms showing the distribution of properties in the database are shown in Figure S5.

### 5.2. Dataset extension for P1 low copy numbers

The dataset above was expanded for the special cases of one or two assemblies in the asymmetric unit and space group P1, to give more data to compare the LESRF from experimental and calculated data. A further condition was added to test case selection for two assemblies in the asymmetric unit: the root-mean-square deviation for all atoms between the two copies of the assembly was restricted to less than 1 Å. We refer to the resulting dataset of P1 with one copy in the asymmetric unit as the “P1-monomer” dataset and with two copies in the asymmetric unit as the “P1-dimer” dataset.

To generate calculated structure factors for the assembly suitable for a SRF, another protocol from the *Phasertng* suite, *Nomad*, was used. Structure factors were calculated for a P1 box with cell dimensions twice the monomer/dimer molecular radius (note that calculated structure factors do not use the unit cell of the crystal). The large unit cell ensures the radius of integration contains only intra-molecular vectors. The calculated structure factors could be used in place of experimental data in *Xtricorder* by ignoring the phase component, reading the σ_A_ values as *D_obs_* and the *E_C_* as *E_eff_*.

## 6. Radius of integration

The SRF (Patterson and LESRF) has a major dependency on the integration radius. If the radius is too small, the function will emphasize secondary structure features such as the symmetry of parallel helices whereas a radius too large will introduce noise from intra-molecular vectors.

How well the intra-molecular vectors are isolated from the inter-molecular vectors by restriction of the radius of integration depends on the sphericity/anisotropy of the atomic distribution of the biological assembly (the macromolecular monomer or macromolecular complex), and the packing/assembly associations in the crystal (Evans & McCoy, 2008). Although the former may be estimated prior to structure solution (see below), the latter cannot.

The integration radius for the SRF is maximally twice the molecular radius, where all vectors are included, but there is no consensus on the appropriate fraction/multiple (less than 2) to use to optimize the signal in the SRF.

### 6.1. Estimation of the molecular radius

In *Xtricorder* the user has three options for providing information about the molecular radius.

If a model is provided, then the molecular radius is taken as the radius that encompasses 90% of the atoms in the model, *R_M90_*. By taking the radius encompassing 90% of the atoms, volume added by extended N-and C-termini and other low-density/not-compact components of structure at the surface are excluded, since these will not contribute significantly to the Patterson density.

Alternatively, if the molecular weight is provided (or the sequence is provided, from which the molecular weight can be calculated) then the molecular radius is estimated as

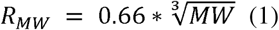

where *R_MW_*is the minimum radius in Ångstroms that can encompass molecular weight *MW* in Daltons (Erickson, 2009).

Lastly, the user may input a molecular radius explicitly (*R_i_*). The fallback is a default molecular radius of 18 Å, which corresponds with MW of 30 kDa via equation (1), 30 kDa being the average molecular weight of a protein domain, likely the smallest component of any assembly crystallized.

The model, sequence or user derived estimates of the molecular radii (*R_MW_, R_M90_, R_i_*) are then adjusted by a scale factor (*k*) of maximum value 2 to give the SRF radius of integration (*b*). Optimization of *k* is discussed below.

Note that *Xtricorder* does not require any information about the assembly radius/model/sequence for the data analysis steps, although it is used for the cell content analysis step if provided.

### 6.2. Constraint on radius of integration

There is a limit on the computational stability of the SRF, with the function being unstable beyond a spherical Bessel function *L_max_* of 100. The *L_max_* of the spherical Bessel functions in turn depends on the radius of integration (*b*) and the maximum resolution (*d_min_*) of the data through the formula,

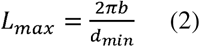

There is thus a limit on the radial resolution of the analysis, either though the data resolution or maximal vector length, the latter because the maximal vector length determines the sensitivity of a change in atomic positions (at the vector-defined molecular boundary) to a given rotation angle.

In *Xtricorder*, the default resolution for the SRF is 3 Å, so the maximal integration radius (*b_max_*) is 48 Å. Using an integration radius equal to the molecular radius (*k = 1*), this corresponds with a protein of molecular weight approximately 170 kDa.

Anisotropic diffraction limits cannot be imposed in the SRF. Since data in the weakly diffracting direction will not contribute signal to the SRF, anisotropic data should be truncated isotropically at a suitably lower resolution limit when the radius of integration is constrained by the limit on *L_max_*.

### 6.3. Theoretical Cumulative Patterson Density

Theoretical plots of the density of Patterson peaks as a function of a fraction/multiple of the molecular radius are shown in Figure 1. Even with very different molecular shapes, ranging from extended structures to compact folds, the plot of the density of Patterson vectors varies most between folds at less than 0.6 times the molecular radius, converging to 70% of the maximal density regardless of fold at around 0.8 times the molecular radius and reaching 95% of the maximal density by 1.4 times the molecular radius.

**Figure 1.**
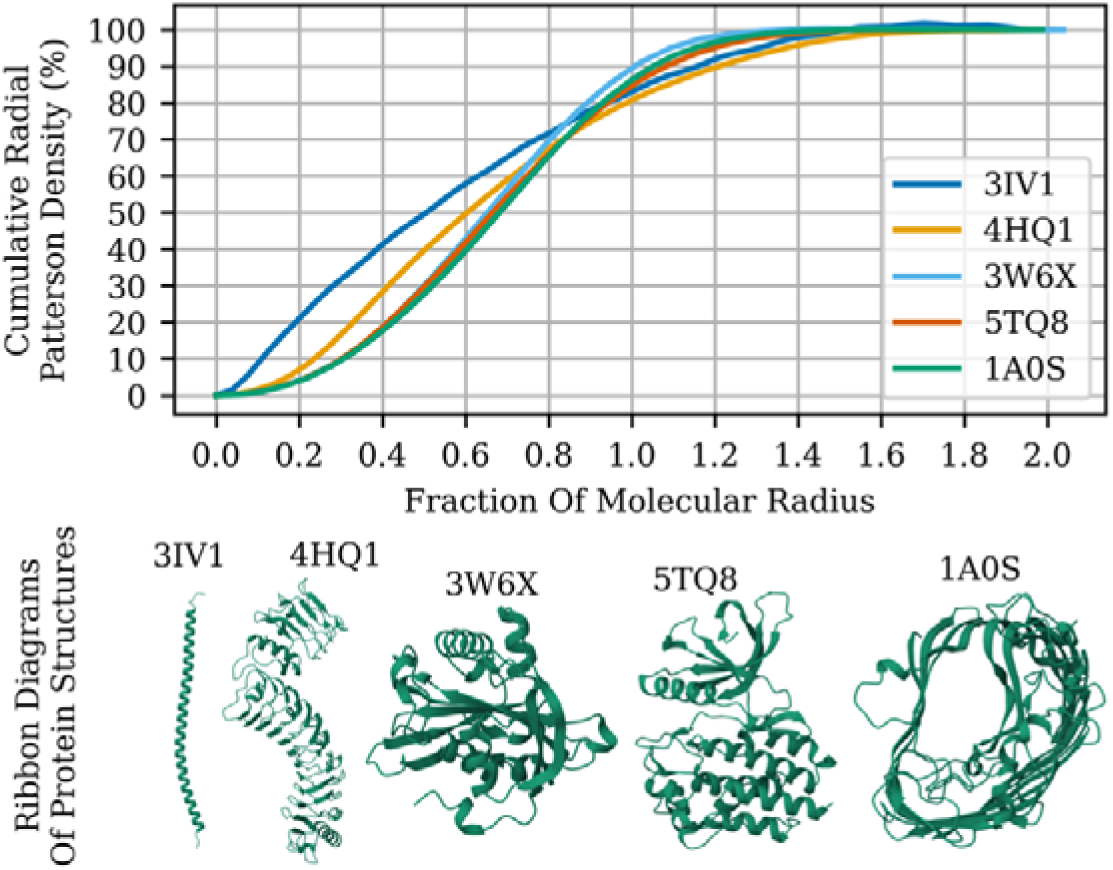
Spatial distribution of Patterson density for representative protein structures. The cumulative radial Patterson density is plotted as a function of the fractional molecular radius for five protein structures chosen to differ markedly in fold: 3IVI is an extended α-helix extracted from a coiled coil, 4HQ1 is an extended leucine-rich repeat with two solenoids, 3W6X is a highly globular protein, 5TQ8 is a bi-lobal kinase and 1A0S is a porin. For each structure, the atomic model was centred at the origin, placed in a large cubic P1 unit cell, and used to compute a map. The squared amplitudes of the Fourier coefficients were spherically averaged in concentric shells to yield the radial distribution of Patterson density. The cumulative sum of shell contributions, normalized to 100%, quantifies the proportion of total Patterson signal within a given radius. Ribbon diagrams of the protein structures are shown for visual reference, with PDB codes indicated. Each image was rendered using *Mol* Viewer* (Sehnal *et al*., 2021).

### 6.4. Optimization of radius of integration

The relationship between the radius of integration and the LESRF results were probed using the ‘P1-monomer’ and ‘P1-dimer’ datasets. The ‘P1-monomer’ dataset was more suited to probing the noise-like features of the LESRF due to crystal packing and pseudo-symmetries, and the ‘P1-dimer’ dataset was more suited to probing the signal-like features in the LESRF. Signal in ‘P1-monomer’ cases could only arise from internal domain duplications (if present) or extended secondary structural elements, particularly helices, while for the ‘P1-dimer’ data cases it was the symmetry mapping one copy to the other.

The SRF was run using molecular radius *R_M90_* with a scaling factor *k* taking 6 values of 0.6, 0.8, 1.0, 1.2 and 1.4. The SRF results for each test case were compared by analysing the correlation between the distributions of top SRF peak heights across cases and performing a regression analysis, as shown in Figure 2. Three metrics – the Pearson correlation coefficient, the silhouette score and the centroid distance –were considered to optimize *k*.

**Figure 2.**
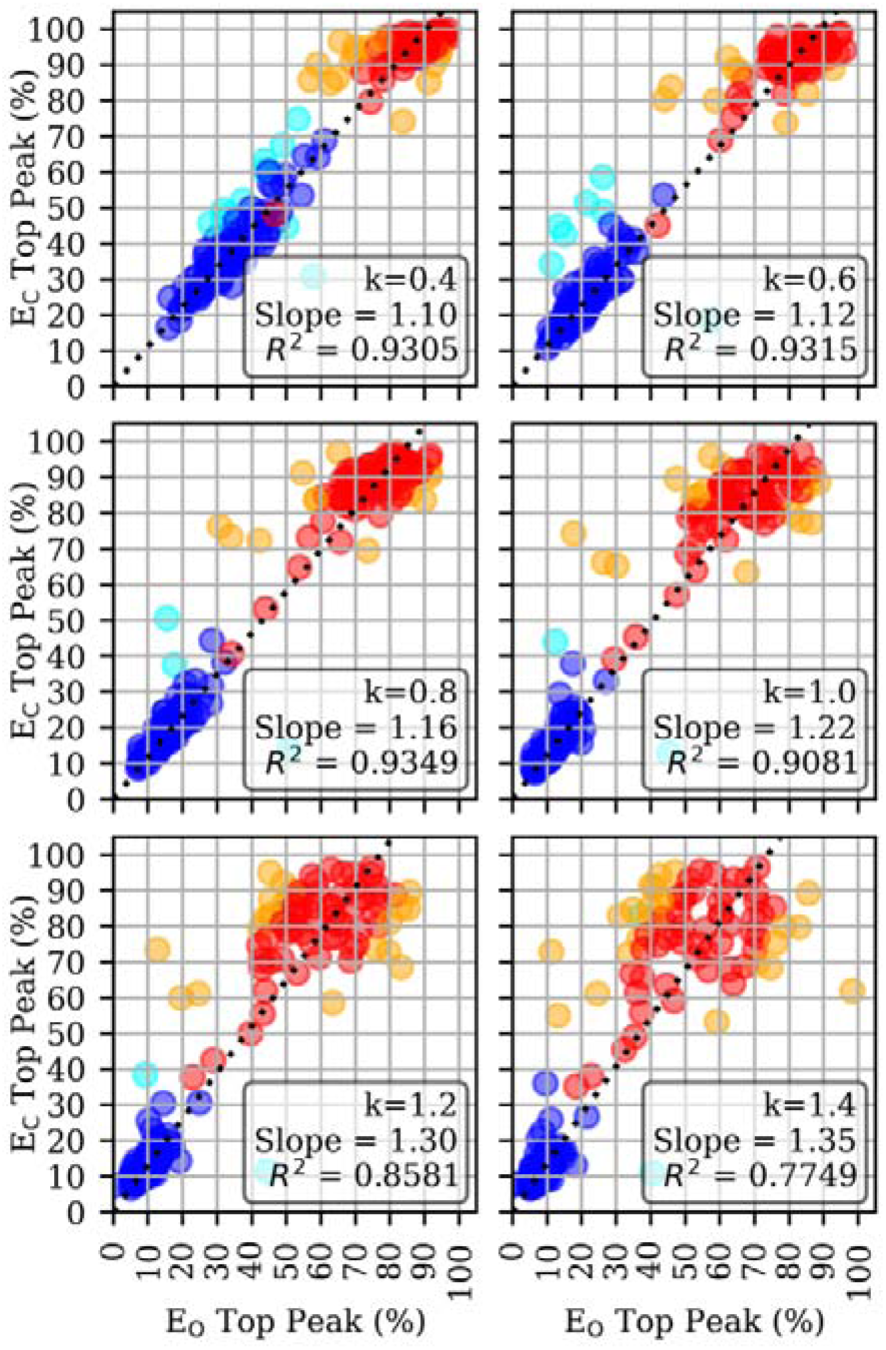
Correlation and clustering analysis of SRF peak prominence across different integration radii. Scatter plots show the relationship between the top SRF peak in calculated versus observed datasets for ‘P1-monomer’ and ‘P2-dimer’ datasets (blue and red, respectively) across six *k* values where *k* is the scale factor for the molecular radius *R_M90_* (see text). Each subplot (a–f) corresponds to a different *k* value (*k* in range 0.4 to 1.4 in steps of 0.2). A dashed black line indicates the least-squares regression line constrained through the origin. Outliers (points with residuals >2σ from the forced regression line) are shown in cyan (‘P1-monomer’) or orange (‘P2-dimer’).

Pearson correlation coefficients between the SRF highest peak height per case for the observed and calculated data per *k* showed a maximum at *k* = 0.8 (Figure 3).

**Figure 3.**
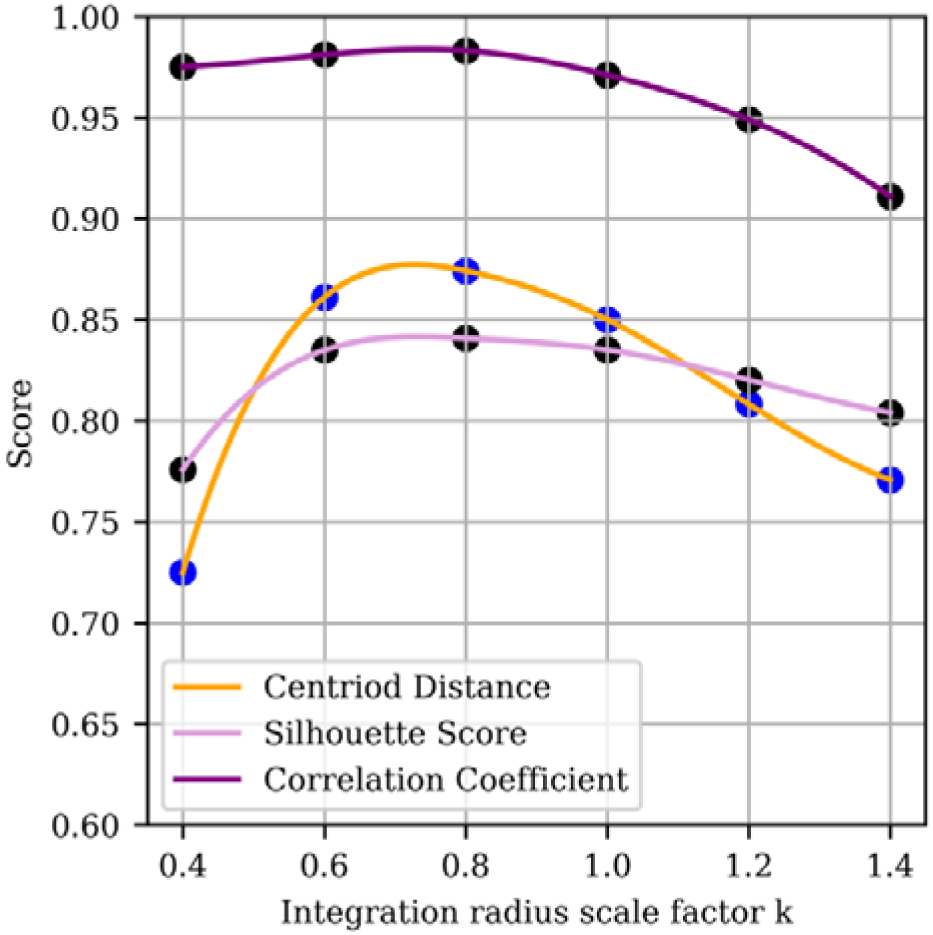
Comparison of ‘P1-monomer’ and ‘P1-dimer’ class separation metrics as a function of integration radius expressed as the scale factor k of the molecular radius R_M90_. Data as in Figure 2. Three scoring criteria were plotted for varying values of *k* from 0.4 to 1.4: the correlation coefficient between observed and calculated peak prominence (purple), centroid distance between cluster centres (orange), and silhouette score (plum) as described in the text. Curves were generated by cubic spline interpolation of six empirical values (shown as scatter points). The three metrics have no units.

The silhouette score for a point is calculated by taking the difference of its average distance to all other points in the same cluster (cohesion), and the average distance to all points in the other cluster (separation). It takes values between 1.0 (cohesive and separated), and -1.0 (suggesting misclassification, were classification uncertain), with values of 0.0 for points on the boundary. The overall silhouette score, the average of all individual scores across the dataset (Figure 3), showed a maximum at *k* = 0.8.

The centroid distance is a calculated as the distance between the centroids of the clusters; this metric does not capture the spread of the clusters. The centroid distance scores per *k*, showed a maximum at *k* = 0.8

The three measures agreed in indicating that the optimum was *k* = 0.8. Note that the density of Patterson peaks at *k* = 0.8 also benefits from not being particularly sensitive to molecular shape (Figure 1). This value was used for all later analysis.

### 6.5. Applicability to other SRF implementations

The optimization determined here is relevant to other implementations of the SRF. In *CNS*, *X-PLOR*, and *Polarrfn* the molecular radius is required user input. In *Molrep*, the user can choose instead to input the molecular weight of the protein, from which a radius (method undocumented) is calculated. *Molrep* documentation suggests without attribution that an appropriate integration radius is twice the radius of gyration. Since the radius of gyration for a solid sphere radius *r* is 0.44*r*, the suggested integration radius (for a ‘spherical’ protein) is 0.88*r* which is close to the value of 0.8*r* found to be optimal in this study.

### 6.6. Applicability to CRF

The signal from the CRF is degraded by inaccuracies in the model structure, while the signal from the SRF is enhanced because there is no ‘model’ inaccuracy (only the relatively minor errors in the measurement of experimental data). Conversely, the CRF signal is enhanced by the ability to remove any inter-molecular vectors by calculation of the structure factors in an unphysiological crystal created such that the assemblies in the lattice are separated by over twice their molecular radii, while the SRF signal is degraded by unavoidable inter-molecular vectors (since the crystal forms a connected lattice).

These highly divergent properties of the CRF versus the SRF mean that optimizations of the radius of integration *b* relevant to the SRF will not translate to optimizations of the integration radius for the CRF. Indeed, we have long established that for the CRF it is optimal to include all vectors (Storoni *et al*., 2004).

## 7. Asymmetric unit copy number

We aimed to improve the prior probabilities for the number of copies in the asymmetric unit by inclusion of information from the LESRF in the case of general non-crystallographic symmetry. The data extracted from the PDB was sparse for 11, or more than 12 assemblies in the asymmetric unit. Therefore, our predictions aimed to classify the data into one of 12 classes (1, 2, 3, 4, 5, 6, 7, 8, 9, 10, 12, ’X’) where X was for 11 or more than 12, serving also as a catch-all to accommodate uncertainty. This allowed the prediction to focus its discriminative power on the well-populated classes, while still accounting for rare or ambiguous instances without compromising overall accuracy.

### 7.1. Matthews coefficient

Prior probabilities for the number of assemblies present in the asymmetric unit have long been established from the Matthews coefficient (Matthews, 1968). Most non-viral protein structures have a protein content within 20%–70% (Kantardjieff and Rupp, 2003).

The Matthews coefficient is a stringent prior when the molecular volume to asymmetric unit volume ratio is high. It is diagnostic for one copy if one copy has a protein content over 50%. If the hypothetical presence of one copy in the asymmetric unit gives less than 50% protein, in practice the protein content for one copy needs to be less than 35% to also allow for two copies within the 20%– 70% range, and less than 23% to also allow for three copies. The Matthews coefficient becomes progressively less informative about asymmetric unit copy numbers as the molecular volume to asymmetric unit volume decreases.

The solvent content of protein crystals is known to be correlated with the maximal diffraction resolution of the crystal, since tighter packing leads to higher order in the lattice. Different space groups have been investigated for correlation with different average solvent contents, with it having been observed that more frequent space groups have lower average solvent contents (Andersson and Hovmöller, 2000). However, a later study indicated that the important aspect of differences in solvent content between space groups was the symmetry L-value, the number of independent parameters for describing the unit cell (Wukovitz and Yeates, 1995) and the average Vm ranges from 2.40 for triclinic lattices to 3.44 for cubic lattices (Chruszcz *et al*., 2008).

In the special case of commensurate tNCS, the number of copies in the asymmetric unit can be deduced from the tNCS vectors. This is most easily done when the vectors are expressed in fractional coordinates. For example, a single tNCS vector at (½,0,0) implies a multiple of two in the asymmetric unit, and the set of vectors (¼,0,0), (½,0,0) and (¾,0,0) imply a multiple of four in the asymmetric unit.

### 7.2. Pilot Study with Decision Tree

Our previous work on tNCS established criteria for the recognition of tNCS from the Patterson. In summary, a native Patterson peak of 16% identified a significant tNCS, where significance was defined by the need to correct the intensities for the tNCS-associated systematic intensity modulations before use in the maximum likelihood functions for molecular replacement (Caballero *et al*., 2020). We wished to establish a similar criterion for the distinguishing between the presence and absence of general NCS using the SRF.

As a pilot study, we selected from our dataset monomer and dimer cases for all space groups. Unlike the study for the optimization of integration radius, we included all cases regardless of the root-mean-square deviation between the copies in the asymmetric unit (for cases of two in the asymmetric unit).

Histograms of the distribution of top non-origin peak heights as a percentage of the origin (Figure 4), were analysed to find the Kolmogorov-Smirnov (KS) threshold, the point on the cumulative distribution curves where the separation between classes is maximized, which is the decision boundary where the true positive rate and false positive rate differ the most.

**Figure 4.**
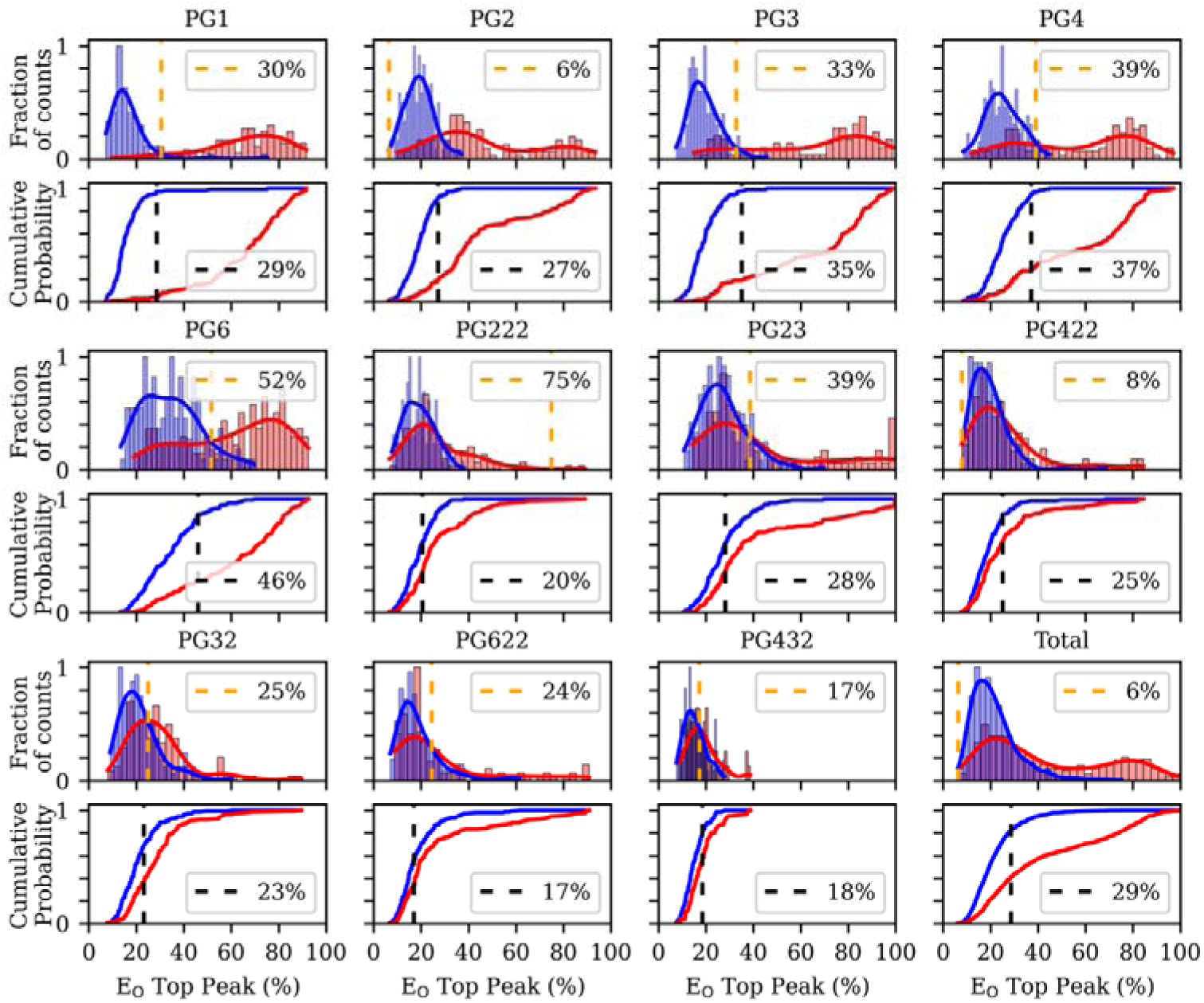
Distributions of SRF peak height values by symmetry group and oligomeric state. Each symmetry group (PG1 to PG432, plus a combined “Total”) is represented by a pair of vertically stacked plots: a density-normalized histogram (top) and a cumulative distribution function (ECDF; bottom). Blue and red indicate data from ‘P1-monomer’ and ‘P1-dimer’ datasets respectively. In the histograms, the shaded bars show the relative frequency distributions of SRF peak height (%) values with overlaid kernel density estimates (KDEs), each normalized to a maximum of 1. An orange dashed line marks the KDE-based threshold at which the densities of the two classes are most similar. In the ECDF plots, a black dashed line indicates the value at which the absolute difference between monomer and dimer cumulative distributions is greatest (Kolmogorov–Smirnov statistic). Axes are shared within rows; x-axis labels appear only on the lower subplots, and y-axis labels only on the leftmost panels.

The ability to distinguish one from two in the asymmetric unit (two classes for categorization) became progressively more difficult as the number of crystallographic space group operations increased. For more than six symmetry operations, the SRF top peak height distributions for the two classes overlapped extensively. No peak height cutoff gave good discrimination between the two classes.

Attempts to add other criteria to the classification, from amongst the many in the database associated with the dataset, via a decision tree, failed to find any combination of criteria that improved the ability to distinguish classes (data not shown).

The failure to find a SRF peak height that indicated the presence of general non-crystallographic symmetry even in this simple pilot study indicated that a decision tree approach would not yield helpful insights when extended to higher order non-crystallographic symmetry.

### 7.3. Circular, Dihedral, Octahedral and Icosahedral symmetry

The relationship between the number of copies of the assembly in the asymmetric unit components and the presence of circular, dihedral, octahedral and icosahedral symmetry in the SRF in our dataset is shown in the series of heatmaps in Figure 5. The data is presented in a stacked bar chart in Figure S6.

**Figure 5.**
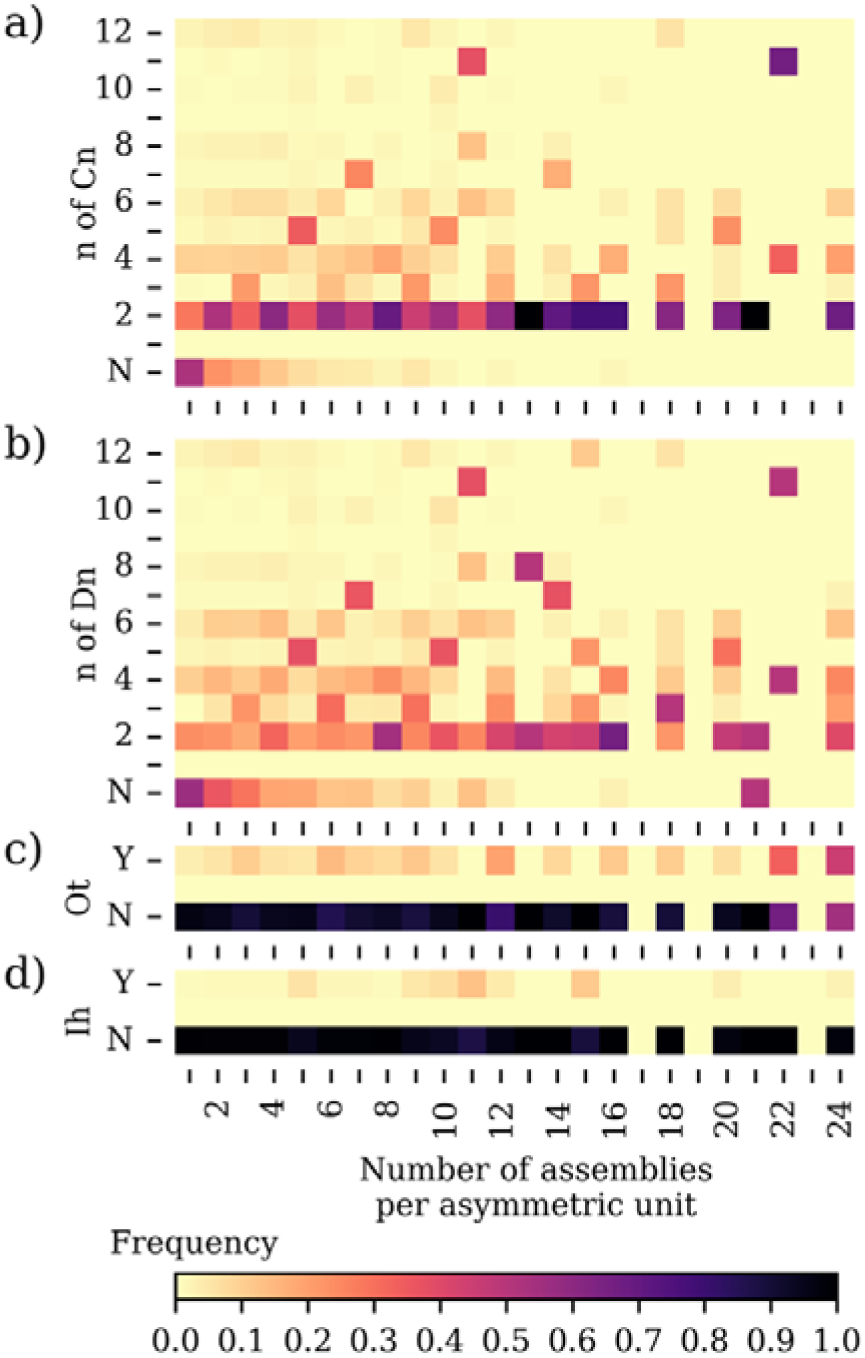
Normalized frequency heatmaps of SRF symmetry indicators by number of assemblies in the asymmetric unit. Each panel shows a 2D histogram of symmetry-related metrics (y-axis) as a function of the number of assemblies per asymmetric unit (x-axis) a) *n* of the point group *C_n_* (y-axis), for most prominent peak over 20% of the origin peak with ‘N’ indicating no peak over 20% of the origin (b) *n* of the point group *D_n_* (y-axis), for most prominent peak over 20% of the origin peak with ‘N’ indicating no peak over 20% of the origin (c) presence of an octahedral symmetry with all peaks for the set of symmetry operators over 20% of the origin (d) presence of an octahedral symmetry with all peaks for the set of symmetry operators over 20% of the origin. Colour intensity indicates the relative frequency (0 to 1) within a given column.

The most common SRF symmetry is C_2_ regardless of the number of copies in the asymmetric unit. The presence of C_2_ symmetry therefore contains little information about the contents of the asymmetric unit. The presence of odd *n* integer C_n_ symmetry as the top peak in the SRF, for example C_5_ C_7_ or C_11_, is much more informative about the number of copies in the asymmetric unit.

Where C_2_ symmetry is present, it can also be a subgroup of higher order oligomers, for example tetramers that are dimers of dimers, with D_2_ symmetry. Where C_n_ symmetries are present as a subgroup of D_n_ symmetry, they may be considered significant even if the C_n_ peak is not the highest in the SRF: commonly it is the associated perpendicular C_2_ symmetry that has highest SRF peak. The presence of odd integer D_n_ symmetries, for example D_5_ D_7_ or D_11_, can be diagnostic for the number of copies in the asymmetric unit even if the associated C_n_ is not the highest SRF peak.

Octahedral symmetry (O_t_), with 24 positions related by rotational symmetry, is rarely found in the SRF in our dataset, except in the case of 24 copies of the assembly in the asymmetric unit.

Icosahedral symmetry (I_h_), with 60 positions related by rotation symmetry, was found at very low incidence in the SRF in our dataset, as our dataset did not include (icosahedral) virus structures.

### 7.4. Methods for Machine Learning

The differences in C_n_, D_n_, O_t_ and I_h_ symmetry detected in the SRF between different asymmetric unit copy numbers indicates that there is information in the SRF that could be interpreted by machine learning (Figure 5, Figure S6).

The data were organized into a three tiered directory structure to facilitate reproducible loading and splitting. At the top level, samples were divided into “train,” “validate” and “test” sets with an approximate 80 %/10 %/10 % split, stratified to balance SGPG distribution as far as possible. Classes for only one or two copies in the asymmetric unit were further restricted to avoid vast overrepresentation (Table 1). Within each split, second level subdirectories were named for the integer number of biological assemblies per asymmetric unit, and third level subdirectories were named for individual PDB identifiers. Each such PDB folder held exactly thirteen 112×112 pixel grayscale images in portable network graphics (PNG) format, representing successive slices of the self rotation function as described above, with the first layer containing the Matthews coefficient encoding. Classes with fewer than 100 samples were pooled into a single ‘X’ category, serving both to stabilize training and to capture out of distribution cases.

**Table 1.** Number of Test Cases for Classification Training, Validation, and Testing by Assemblies per ASU. This table presents the distribution of test cases across the training, validation, and testing sets for each class. The number of available datasets in the PDB restricts the number of samples for some classes. The training, validation, and testing sets are used to train, tune hyperparameters, and evaluate the performance of the TensorFlow CNN model.

| assemblies per<br>asymmetric unit | Number in set<br>'train' | Number in set<br>'validate' | Number in set<br>'test' |
| --- | --- | --- | --- |
| 1 | 596 | 60 | 976 |
| 2 | 616 | 60 | 957 |
| 3 | 664 | 68 | 325 |
| 4 | 644 | 62 | 603 |
| 5 | 778 | 142 | 152 |
| 6 | 740 | 80 | 117 |
| 7 | 123 | 8 | 11 |
| 8 | 564 | 67 | 77 |
| 9 | 149 | 26 | 43 |
| 10 | 196 | 21 | 24 |
| 12 | 378 | 56 | 45 |
| 99 | 244 | 27 | 30 |
| total | 5692 | 677 | 3360 |

Images were streamed on-the-fly using a custom generator that loads each of eight greyscale PNG files per sample, stacks them along the depth axis to produce a 3D tensor of shape (112, 112, 8), and pairs each tensor with a one-hot encoded class label based on directory structure. This generator was wrapped in a TensorFlow Dataset pipeline configured for high-throughput execution on a 120-core CPU node under the ‘CentralStorageStrategy’. The pipeline applies per-image standardization, shuffling for training and validation sets, batching (32 samples per batch), and asynchronous prefetching to maximize efficiency.

The classification model was implemented in ‘Keras’ using the Sequential API as a pruned 3D convolutional neural network. It comprises three ‘Conv3D’ layers with increasing filter depths (32, 64,128), all using 3×3×3 kernels, ‘Re U’ activations, and ’same’ padding. Each ‘Conv3D’ layer is followed by MaxPooling3D that pools over the two spatial dimensions but preserves depth. The resulting 3D feature volume is flattened and passed through a 256-unit dense layer with ‘Re U’ activation and a dropout rate of 0.5, before being projected by a final dense layer with ‘softmax’ activation onto the 12 output classes.

To reduce model size and improve efficiency, layer-wise magnitude-based pruning was applied to the convolutional layers using TensorFlow Model Optimization Toolkit. A polynomial pruning schedule progressively increased sparsity from 0% to 50% over the course of training. The network was trained for up to 25 epochs using the Adam optimizer with an initial learning rate of 1×10, subject to a custom learning rate scheduler that reduced the rate at epochs 5, 10, 15, and 20. Early stopping with patience of 5 epochs monitored validation loss and restored the best weights. Model selection was based on validation accuracy rather than loss. After training, pruning wrappers were stripped to yield a deployable, compact model, which was saved in HDF5 format (320Mb).

During the evaluation phase, the model’s performance was measured using test data that was not seen during training. Performance was evaluated via overall accuracy, a classification report of precision, recall, f1-score and support, and a confusion matrix.

### 7.5. Control study

As a control, the Matthews coefficient was encoded as a graphic, as above, but with no other information from the SRF (Figure S4). The control served as a baseline for analysing the architecture of the training on the predictions. The convolutional neural network architecture was adapted from the primary study, with the key difference that only a single greyscale image was loaded per sample, and the three convolutional layers were implemented as 2D convolutions (Conv2D) rather than 3D (Conv3D).

### 7.6. MLE-Matthews analysis of asymmetric unit contents

The results of the machine learning are compared with the Matthews coefficient by asymmetric unit copy number in Table 2 and by space group point group in Table 3. Confusion matrices of the training are shown in Figure 6. The overall accuracy for the Matthews model was 0.44, for the MLE-Matthews Control 0.67 and for the MLE-Matthews 0.81

**Table 2.** -F1-scores for predicted assemblies per asymmetric unit (a.s.u.). The table compares F1-scores across three models: the original Matthews model, the MLE-Matthews Control (trained without SRF information), and the MLE-Matthews model (trained with SRF information). Each row corresponds to a specific number of assemblies per asymmetric unit (1–10,12, other), with the final column indicating class support (the number of test samples for each class). The MLE-Matthews model consistently outperforms, particularly for underrepresented or previously poorly classified classes such as 5, 6, 7, and 10. The “other” class refers to test samples with 11 or more than 12 in the asymmetric unit.

| assemblies per<br>asymmetric unit | F1-score<br>Matthews | F1-score<br>MLE-Matthews<br>Control | F1-score<br>MLE-Matthews | support |
| --- | --- | --- | --- | --- |
| 1 | 0.74 | 0.93 | 0.95 | 976 |
| 2 | 0.40 | 0.77 | 0.86 | 957 |
| 3 | 0.16 | 0.47 | 0.65 | 325 |
| 4 | 0.20 | 0.56 | 0.76 | 603 |
| 5 | 0.16 | 0.27 | 0.76 | 152 |
| 6 | 0.16 | 0.40 | 0.55 | 117 |
| 7 | 0.00 | 0.00 | 0.67 | 11 |
| 8 | 0.07 | 0.48 | 0.58 | 77 |
| 9 | 0.00 | 0.00 | 0.27 | 43 |
| 10 | 0.18 | 0.00 | 0.61 | 24 |
| 12 | 0.14 | 0.39 | 0.57 | 45 |
| other | 0.52 | 0.59 | 0.82 | 30 |
| Macro average | 0.23 | 0.41 | 0.81 | 3360 |
| Weighted average | 0.40 | 0.68 | 0.67 | 3360 |
| Accuracy | 0.44 | 0.67 | 0.81 | 3360 |

**Table 3.** -Accuracy predicted assemblies per space group. The table compares accuracy across two models: MLE-Matthews Control (trained without SRF information), and the MLE-Matthews model (trained with SRF information). Each row corresponds to a specific space group point group, with the final column indicating class support (the number of test samples for each class). The results for

| Space Group<br>Point Group | Accuracy<br>MLE-Matthews<br>Control | Accuracy<br>MLE-Matthews | number |
| --- | --- | --- | --- |
| 1 | 0.73 | 0.87 | 274 |
| 2 | 0.74 | 0.86 | 350 |
| 3 | 0.71 | 0.86 | 247 |
| 4 | 0.67 | 0.83 | 218 |
| 6 | 0.62 | 0.77 | 252 |
| 222 | 0.76 | 0.84 | 1013 |
| 23 | 0.61 | 0.78 | 153 |
| 422 | 0.59 | 0.73 | 286 |
| 32 | 0.50 | 0.72 | 266 |
| 622 | 0.49 | 0.73 | 197 |
| 432 | 0.39 | 0.75 | 104 |
| total | 0.67 | 0.81 | 766 |

**Figure 6.**
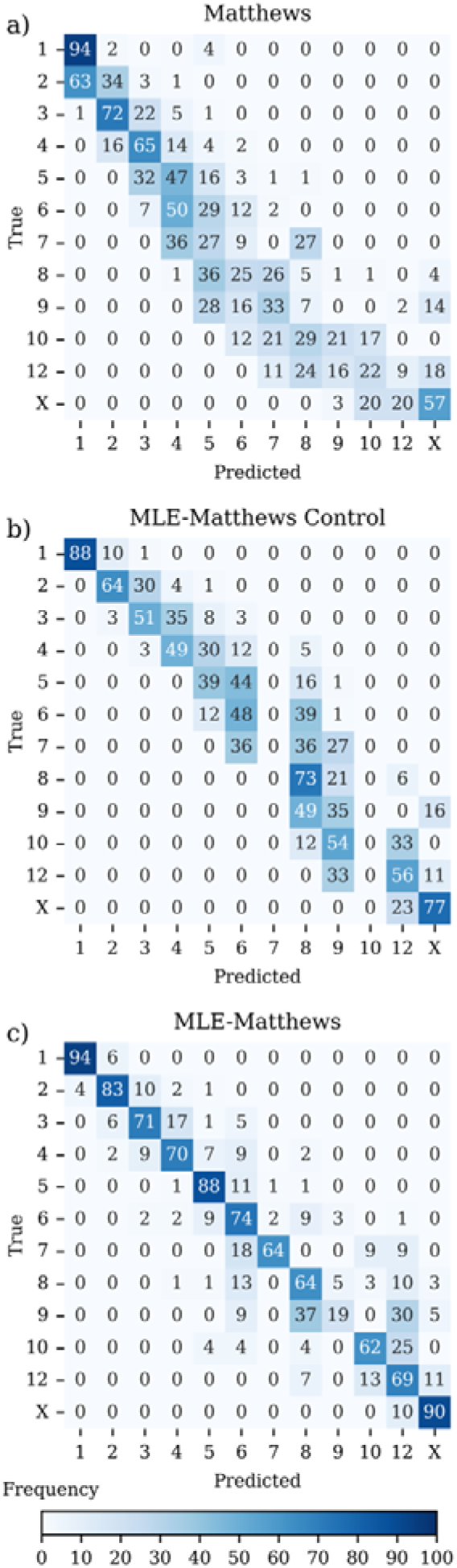
Normalized confusion matrices comparing classification performance of MLE-Matthews against Matthews on test data. The heat maps show per-class prediction distributions for 12 asymmetric unit categories (1-10,12 and ‘X’). Rows represent the true classes, and columns the predicted classes, with values normalized by row to highlight recall performance. Colour intensity corresponds to the percentage of predictions within each true class, with darker shades indicating higher proportions. a) The *Matthews* model displays widespread off-diagonal elements, indicating substantial misclassification, particularly as the number of copies increases. b) The MLE-*Matthews Control* model also displays widespread off-diagonal elements, indicating substantial misclassification. c) The *MLE-Matthews* model has strong diagonal dominance, reflecting accurate and specific classification of structural classes.

The MLE-Matthews Control improves over the Matthews for two reasons. While Matthews predicts unconstrained integer values representing the number of assemblies per asymmetric unit, the MLE-Matthews Control is constrained to select only from the predefined set of allowed classes. This constraint acts as a form of regularization, inherently reducing the chance of erroneous predictions.

Additionally, because the predefined classes reflect the empirical distribution of training data, the constraint introduces an implicit prior: more frequent classes are favoured, biasing the classifier toward commonly observed copy numbers. Consequently, the MLE-Matthews Control achieves higher F1-scores than the Matthews model, despite no new information being added. Performance differences should therefore be interpreted as structural advantages. There is no structural difference between the MLE-Matthews Control and MLE-Matthews.

The MLE-Matthews model consistently outperformed the MLE-Matthews Control. Accuracy was lowest for the higher symmetry space groups, as expected from the pilot study for monomer and dimer.

The inference stage applied a hard limit on the classification dependent on the maximum possible for the asymmetric unit volume. If the predicted class was outside the limit, the highest probability class within limits was used. However, none of the test cases triggered this fallback, which may reflect the effectiveness of the training.

## 8. Discussion

The SRF has sat rather uncomfortably amongst the pantheon of crystallographic data analysis tools. Constantly recommended as a ‘first look’ method, it has never been systematically incorporated into structure solution pipelines, and there are no well-defined rules as to what the crystallographer is to do with what it reveals. The SRF has seemed best interpreted in the rear-view mirror, with the relationship of the SRF peaks to arrangements within the asymmetric unit and crystallographic symmetry operations only discernible *post-hoc*.

A nice example is reported in a CCP4 newsletter article “A Self-Rotation Puzzle” (Cao and Isaacs, 2007), which explores the crystal structure of bovine mitochondrial Peroxiredoxin III (PrxIII), an antioxidant enzyme involved in regulating intracellular hydrogen peroxide levels. Other species of PrxIII had been found to form pentameric rings of dimers (decamers) (Alphey *et al*., 2000). The Matthews coefficient for their novel bovine PrxIII crystal form in space group C2 suggested between 5 and 12 dimers in the asymmetric unit. The SRF showed clear patterns of 2-folds, but did not indicate the expected 5-fold. Rather, there were two 6-fold axes, one 2-fold stronger than the others, and a rotation of 55°. The authors were unable to decipher the SRF *a priori*. Upon phasing by molecular replacement, the crystal structure revealed two half-rings in the asymmetric unit, giving interlocking dodecameric rings in the unit cell — a protein catenane where two closed rings are linked without covalent bonds — which is highly unusual in protein structures. The SRF could then be interpreted as the two 6-folds perpendicular to the planes of the rings, the family of 2-folds perpendicular to the ring axes, and the 55° rotation corresponding to the inclination of the rings (Figure S1, Figure S2.). This remains a difficult case, with MLE-Matthews giving the highest probability (0.89) in the ‘X’ (other/unsure) category.

Attempts have been made to incorporate the symmetries detected in the SRF to enhance the CRF, although the method has not gained traction. The locked rotation function (Tong and Rossmann, 1990), implemented in the program *GLRF* and *Molrep*, aims to take advantage of the symmetry identified in the SRF to average the CRF to reduce noise. A locked rotation function has not been implemented in our software *Phaser* or *Phasertng*, not least because any advantage bestowed by the averaging is diminished by errors in the SRF, and our alternative signal-enhancing method of rescoring CRF identified rotations with a full maximum likelihood rotation function (Read, 2001) is extremely effective, and accounts for errors.

The dominance of C_2_ symmetry in the SRF across all space groups and asymmetric unit contents is not surprising. Proteins commonly evolve dimerization because it is a simple way to evolve stable and versatile function, for example to enable a single protein to act as a switch by toggling its monomeric/dimeric state, or to transmit conformational change for cooperative binding. A weak tendency to self-associate might only require a few mutations to form a stable C_2_ dimer, since the dyad symmetry means that advantageous mutations on one half of the interface are also advantageous at the symmetrical site. The total free energy gain includes enthalpy terms from the formation of electrostatic and hydrogen bonds in the interfaces, and the entropic effect of the exclusion of water, but is reduced by the dimerization itself, which imposes an entropic cost as the two freely diffusing monomers become a dimer. However, if the buried surface area is large enough, typically for a total buried surface of more than 2000 Å², this energetic trade-off tips in favour of dimerization (Chothia and Janin, 1975).

Although the C_2_ dimers present in a crystal lattice may or may not form under physiological conditions, the same symmetrisation of energetic and entropic properties that underly the formation of biological C_2_ dimers also underly the formation of non-physiological C_2_ dimers but facilitated by the high protein concentration environment of the crystallization drop.

Higher order C_n_ oligomers require more complex evolutionary pathways to become stable, since each binding surface is not bilaterally symmetric. While the entropy gains by excluding water from each interface increases with the number of subunits in an oligomer, the entropy cost of association also rises with each additional subunit. Higher-order oligomers thus impose further entropy penalties without necessarily providing proportionally greater stabilization, for the same buried surface area. C_2_ dimers hit the sweet spot, offering a large energetic gain at minimal entropic cost. Higher-order C_n_ oligomers are also difficult to stabilize in a partly associated state and so carry a greater risk of aggregation, and protein insolubility, handicapping any evolutionary advantage. Thus, if C_n_ symmetries with *n*>2 are seen in the SRF, they are more likely to correspond with successfully evolved, biological oligomer states.

The propensity to form dimers also manifests in that higher order C_n_ symmetry commonly present as a subgroup of D_n_ symmetry (Figure 5). This appears to be particularly the case where *n* takes odd values 5, 7, and 11. These symmetries are incorporated in the machine learning, but inspection of the SRF plots by the crystallographer will remain an important step in identifying these more unusual symmetries.

Lower-symmetry space groups (like P1) permit maximal molecular freedom in packing and thus often give tight packing. As symmetries are added, so too are constraints on how the molecules in the lattice must form layered or helical associations. Protein packing around twofold symmetry axes generally involve ‘bump-to-bump’ interactions, while screw axes allow for bump-to-hollow interactions (Filippini and Gavezzotti, 1992): ‘bump-to-bump’ interactions tend to enlarge voids (higher solvent). Empirically, high packing density correlates with higher diffraction resolution. Therefore, differences in average Matthews coefficients for different Bravais lattices (Chruszcz *et al*., 2008) is likely to already be (indirectly) considered for the purposes of asymmetric unit estimation through the resolution dependence of the Matthews probability estimates (Kantardjieff and Rupp, 2003; Weichenberger and Rupp, 2014). Since the space group is a hypothesis in ‘first look’ analysis, particularly regarding the presence or absence of screw axes along any given axis, we have chosen not to incorporate the annotation of the screw symmetries of the refined structure’s space group into our training.

Incorporating information about oligomeric state into structure prediction algorithms can enhance the accuracy of the resulting models, particularly at the interfaces between subunits, which are often highly conserved (Jumper *et al*., 2021). Predicting a protein in the monomeric state when it naturally forms an oligomer may lead to an incorrect fold, because the algorithm may want to place hydrophobic amino acids that should be solvent-accessible in the hydrophobic interior. Tools like AlphaFold-Multimer (Evans et al., 2022; Bryant and Noé, 2024) can leverage known oligomeric states to guide the prediction of both intra- and inter-subunit arrangements, providing more biologically relevant structural models. However, there is currently no way of specifying the point group of the oligomer in AI-generated structure prediction resources, so that although it cannot act as a constraint on prediction, it can be used to select more probable candidates from multiple predictions with different seeds.

The machine learning enhanced (MLE-Matthews) method developed here is the latest in a series of enhancement to the original Matthews coefficients (Matthews, 1968; Kantardjieff and Rupp, 2003; Weichenberger and Rupp, 2014). These studies recalibrated the coefficient over thousands of datasets versus the original few hundred, added separate estimates for protein-nucleic acid complexes, and importantly, included the maximum resolution of the data in the estimate. These enhancements are implicitly included in the MLE-Matthews coefficient through being encoded in the first image.

The new approach integrates features extracted from diffraction data via the LESRF. Our exploration of the peak heights of the LESRF indicated that although the LESRF is not always interpretable *a priori*, on occasions it can be, and this information can be leveraged to increase the likelihood of correctly predicting certain asymmetric unit compositions. The increase in overall accuracy from 0.44 for the Matthew estimation, to 0.67 for the MLE-Matthews Control and to 0.81 for MLE-Matthews is gained mostly from improvements in predictions for higher asymmetric unit copy numbers, and lower symmetry space groups, including space groups P2_1_2_1_2_1_, P2_1_ and C2, space groups that together account for 50% of protein crystal structures (Gaur, 2021): these are also the most useful set of circumstances. As such, this tool offers crystallographers an improved starting point for molecular replacement or experimental phasing, potentially reducing trial-and-error cycles and expediting structure solution.

The MLE-Matthews method is incorporated into *Xtricorder* and the wider *Phasertng* software packages. The results should be used in conjunction with inspection of the SRF plots and tNCS analysis, to corroborate how MLE-Matthews is drawing out likely asymmetric unit compositions. Only inspection of the SRF plots will provide the specific information about the direction of rotation axes and their relationship to each other. Any other complementary lines of evidence such as biophysical studies or known oligomerization state of homologues should always be considered for their support of, or challenge to, the computational predictions, especially if being relied on for AI-generated structure prediction and/or crystallographic phasing.

## Acknowledgements

RJR and AJM acknowledge funding from the Biotechnology and Biological Sciences Research Council (UK) grant number BB/Y009398/1

## Data availability

*Xtricorder* and *Nomad* will be made available through the *PHENIX* and *CCP4* software packages.

