## Supplementary Information for "Xtricorder: A likelihood-enhanced self-rotation function and application to a machine-learning enhanced Matthews prediction of asymmetric unit copy number"

Figure S1 *Structure of bovine mitochondrial peroxiredoxin III (PDB ID: 1ZYE* (Cao *et al.*, 2005)*), a protein catenane comprising two interlocked dodecameric toroids.* The assembly crystallizes in space group *C2* and the images show half of the unit cell. The two rings are depicted in orange and teal to emphasize their topological interlinking. (a) View along the crystallographic *x*-axis, highlighting the central cavities of the toroids. (b) View along the *y*-axis, showing the 55° inclination of the rings. Molecular surfaces were rendered using Mol* Viewer (Sehnal *et al.*, 2021).

a)

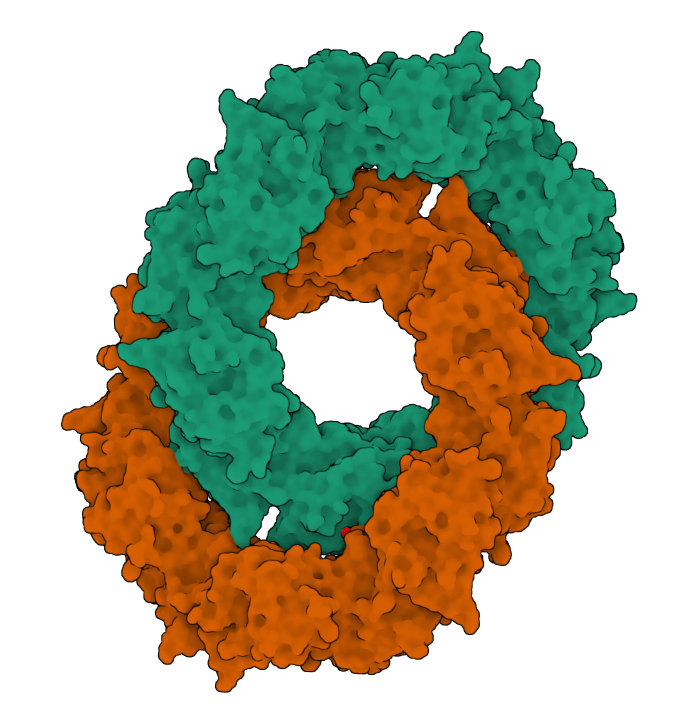

b)

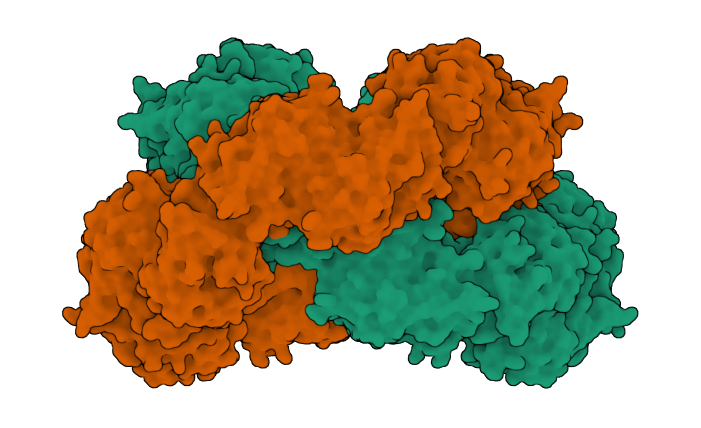

Figure S2 Scatter plots for 1ZYE showing Mercator and stereographic projections of the SRF peak heights. Image is the output pdf, page size A4, from Xtricorder.

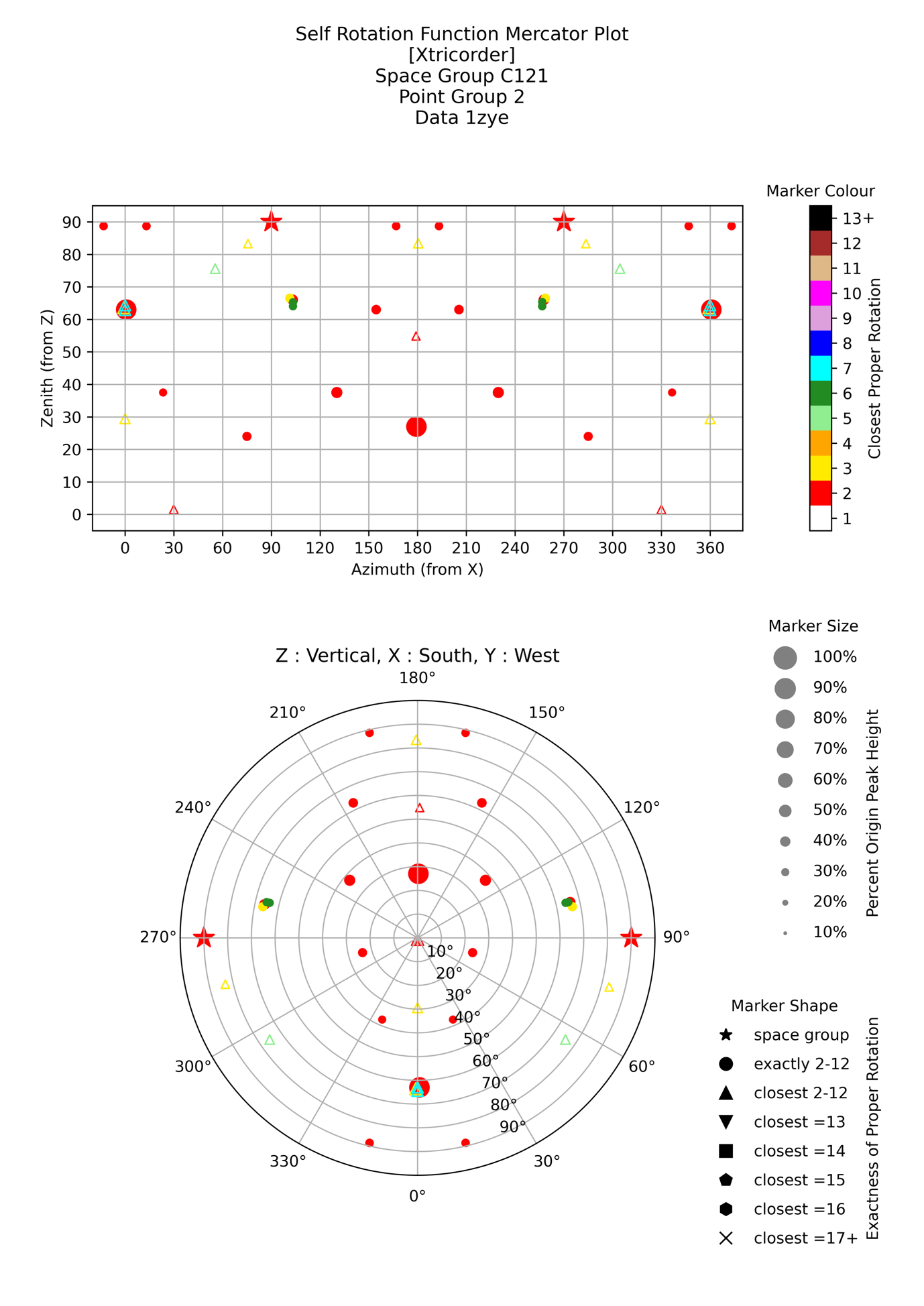

Figure S3 *Self-Rotation Function (SRF) Contour Plots*. Each panel (a–h) shows a 2D contour representation of a SRF section taken at a specific κ‑section, selected to correspond to characteristic n-fold rotational symmetries, with κ ≈ 360°/n for n = 2 to 9 [180°, 120°, 90°, 72°, 60°, 51°, 45°, 40°]. The scatter plot is overlaid.

a)

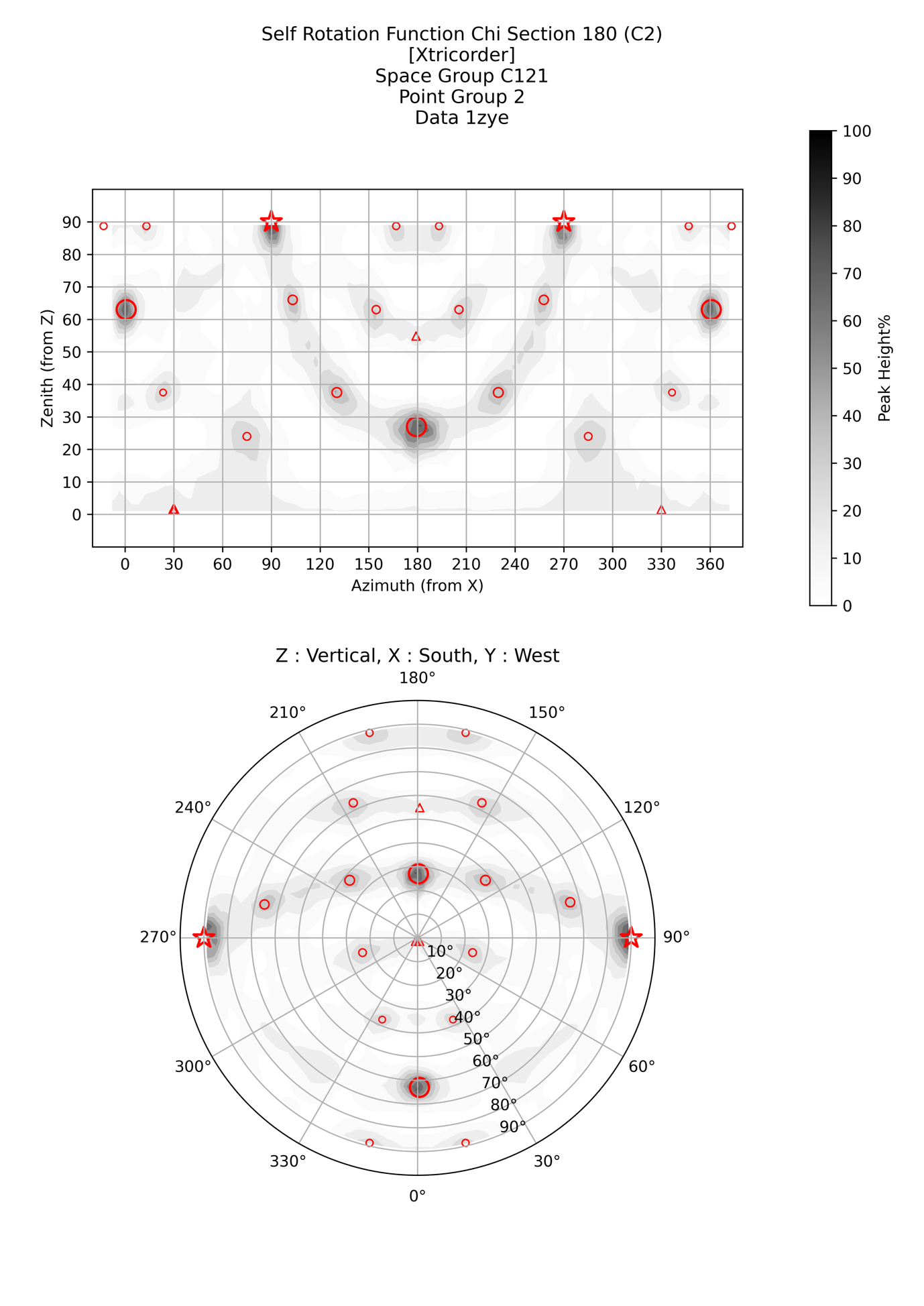

b)

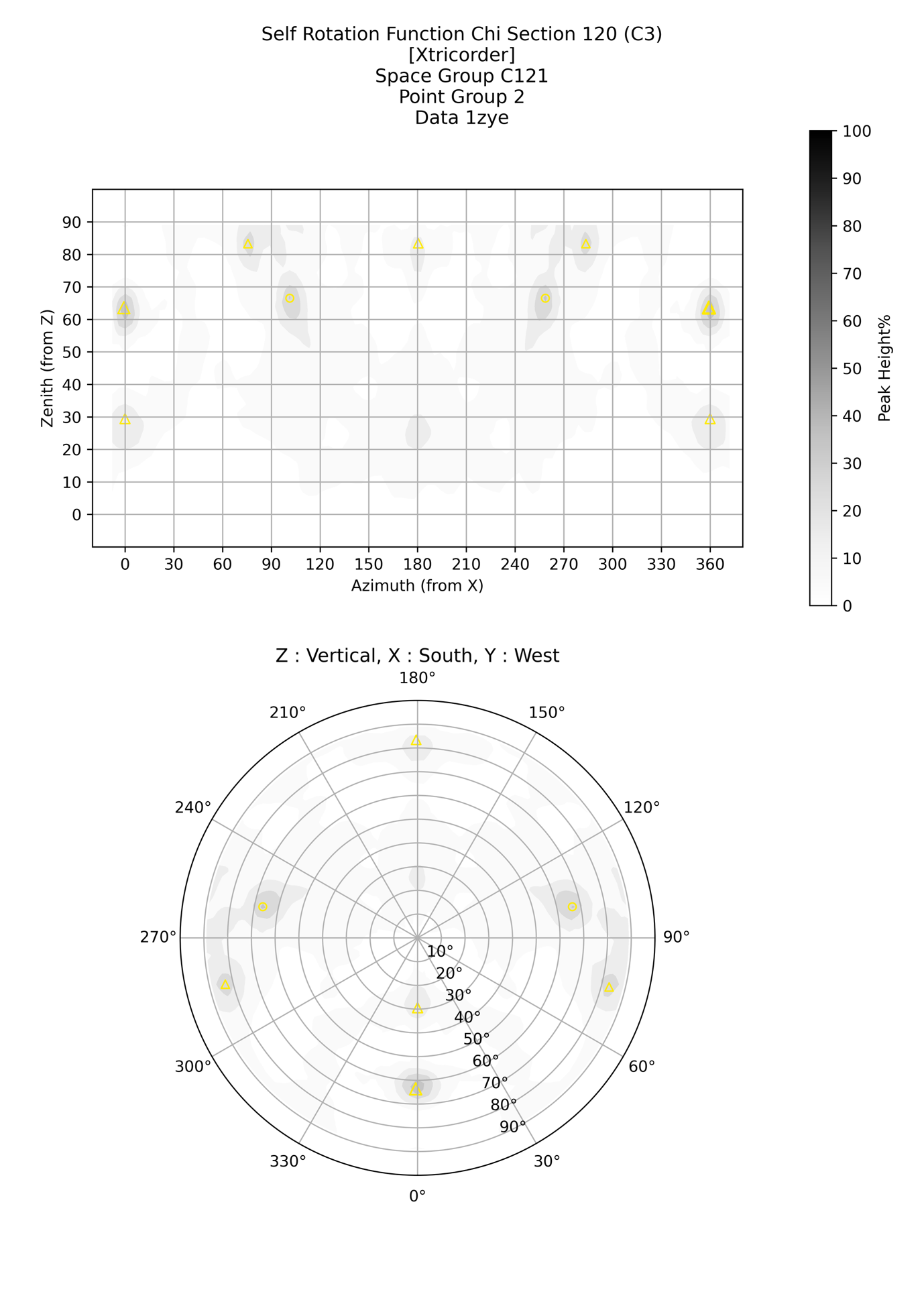

c)

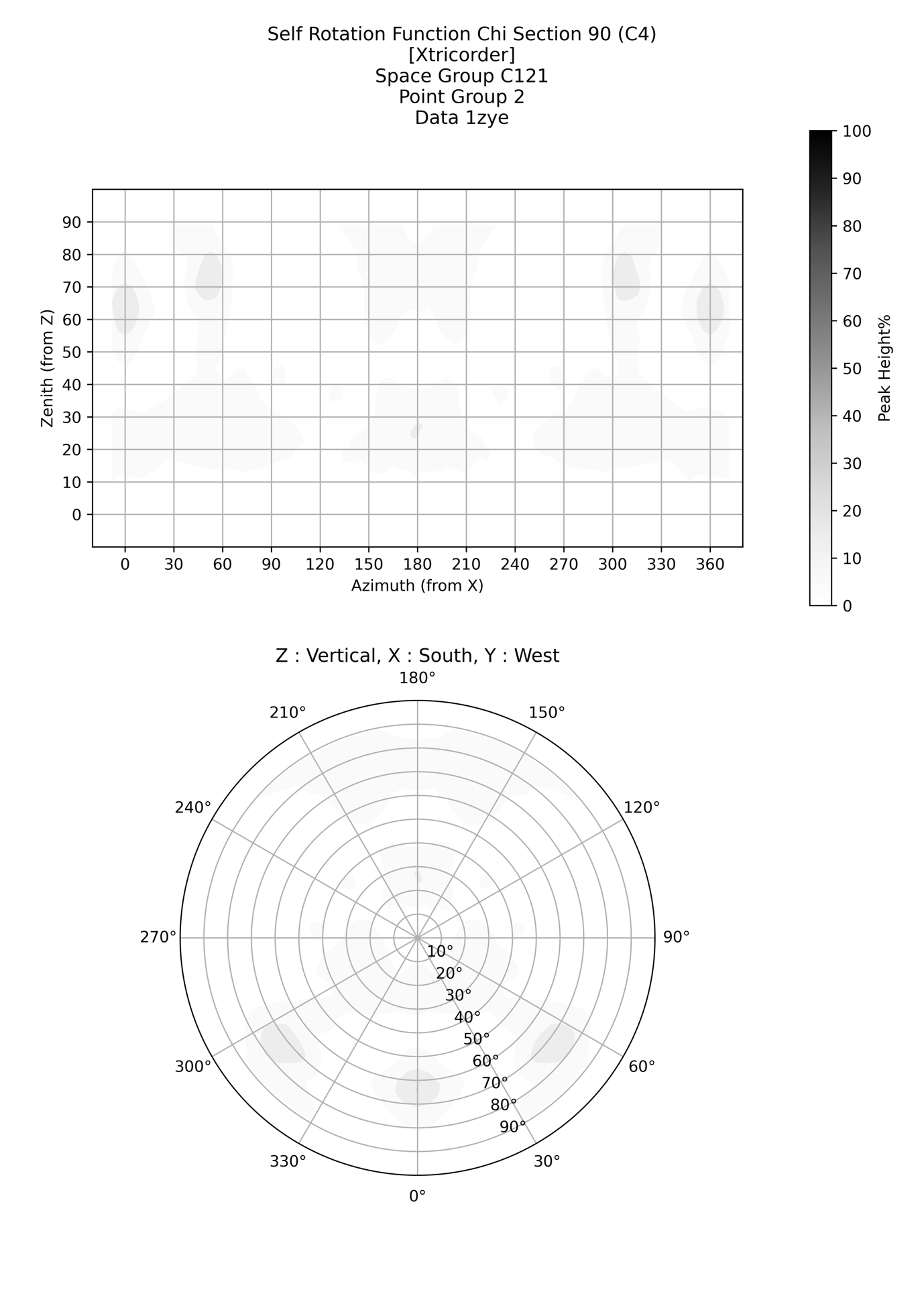

d)

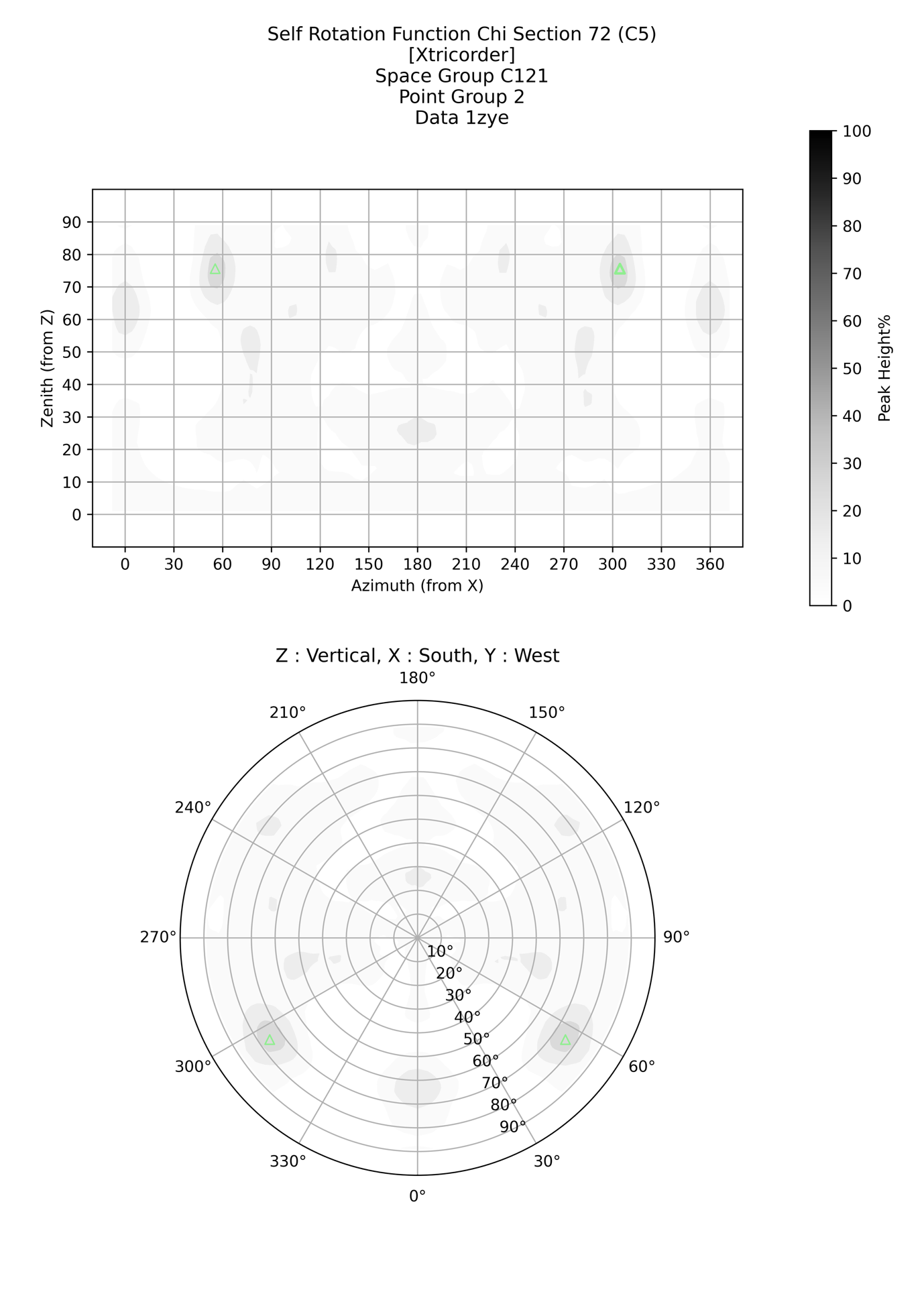

e)

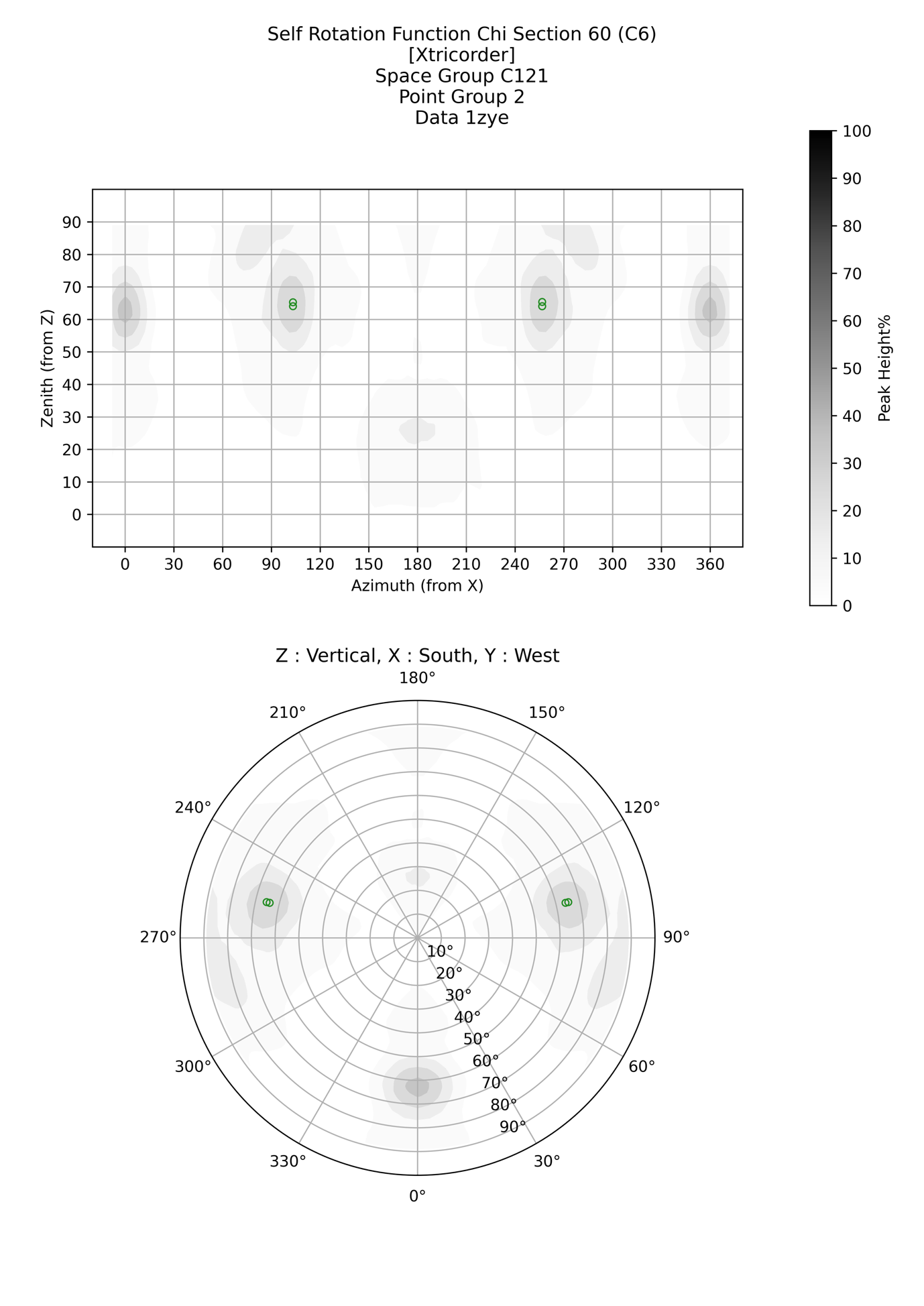

f)

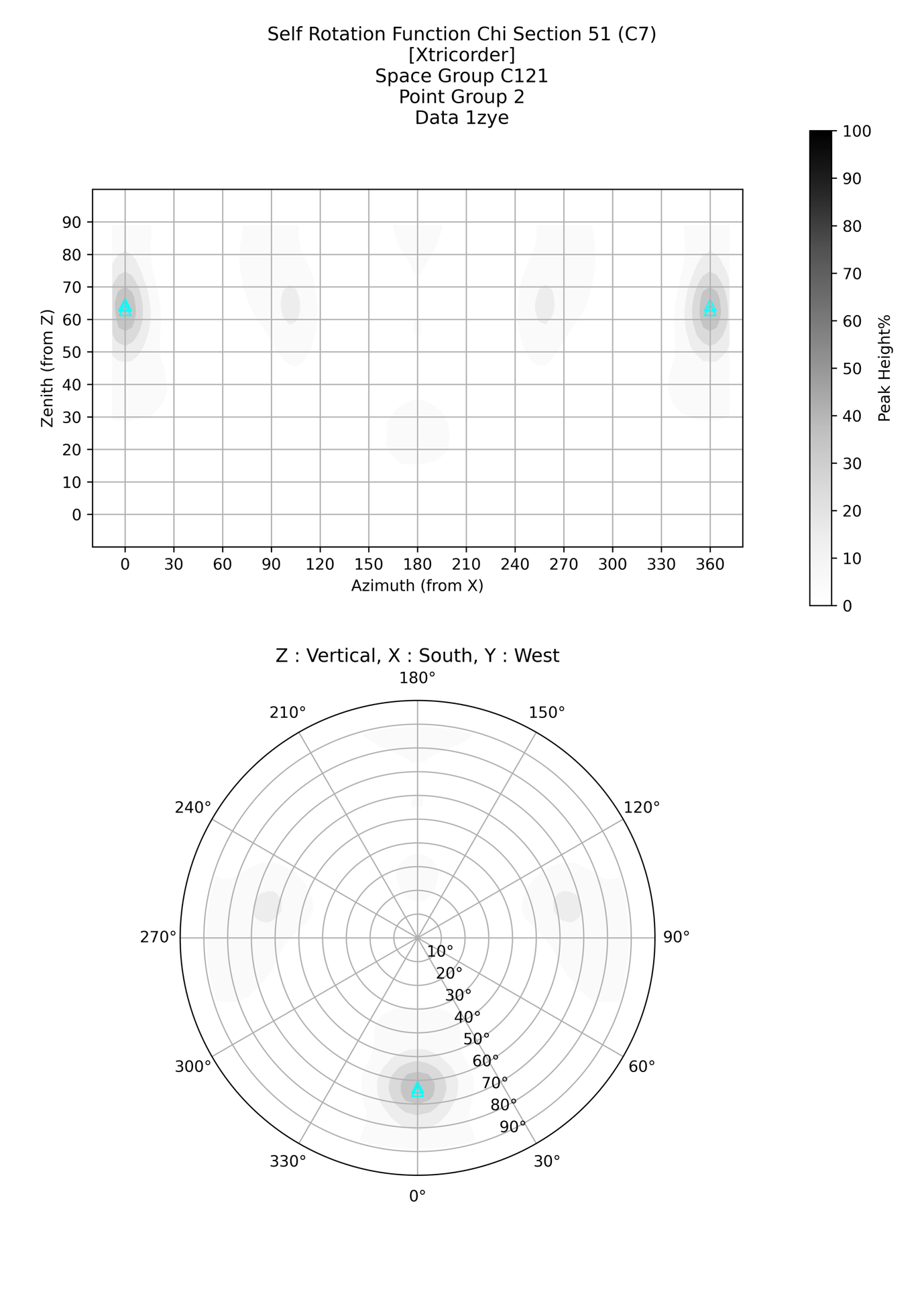

g)

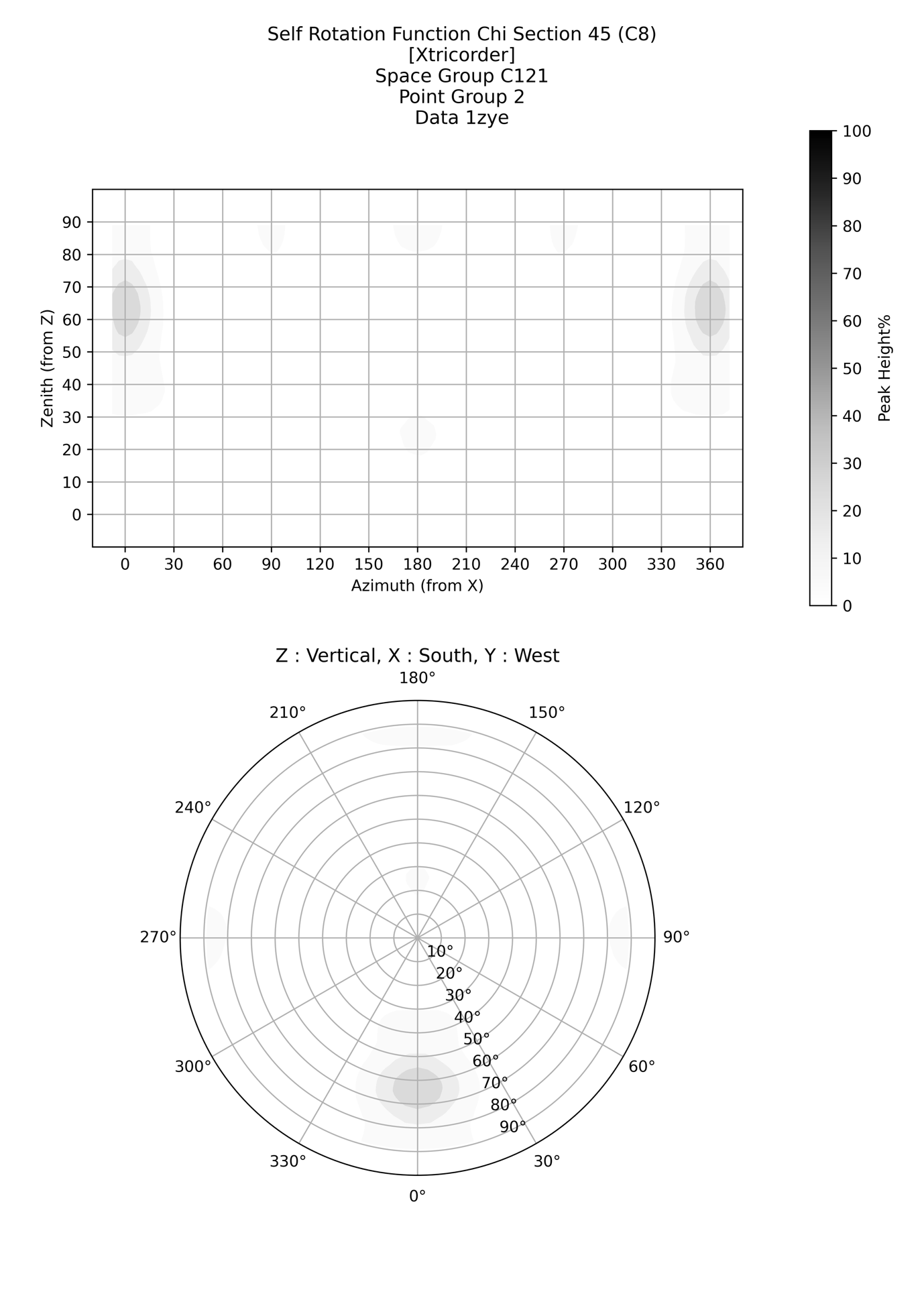

h)

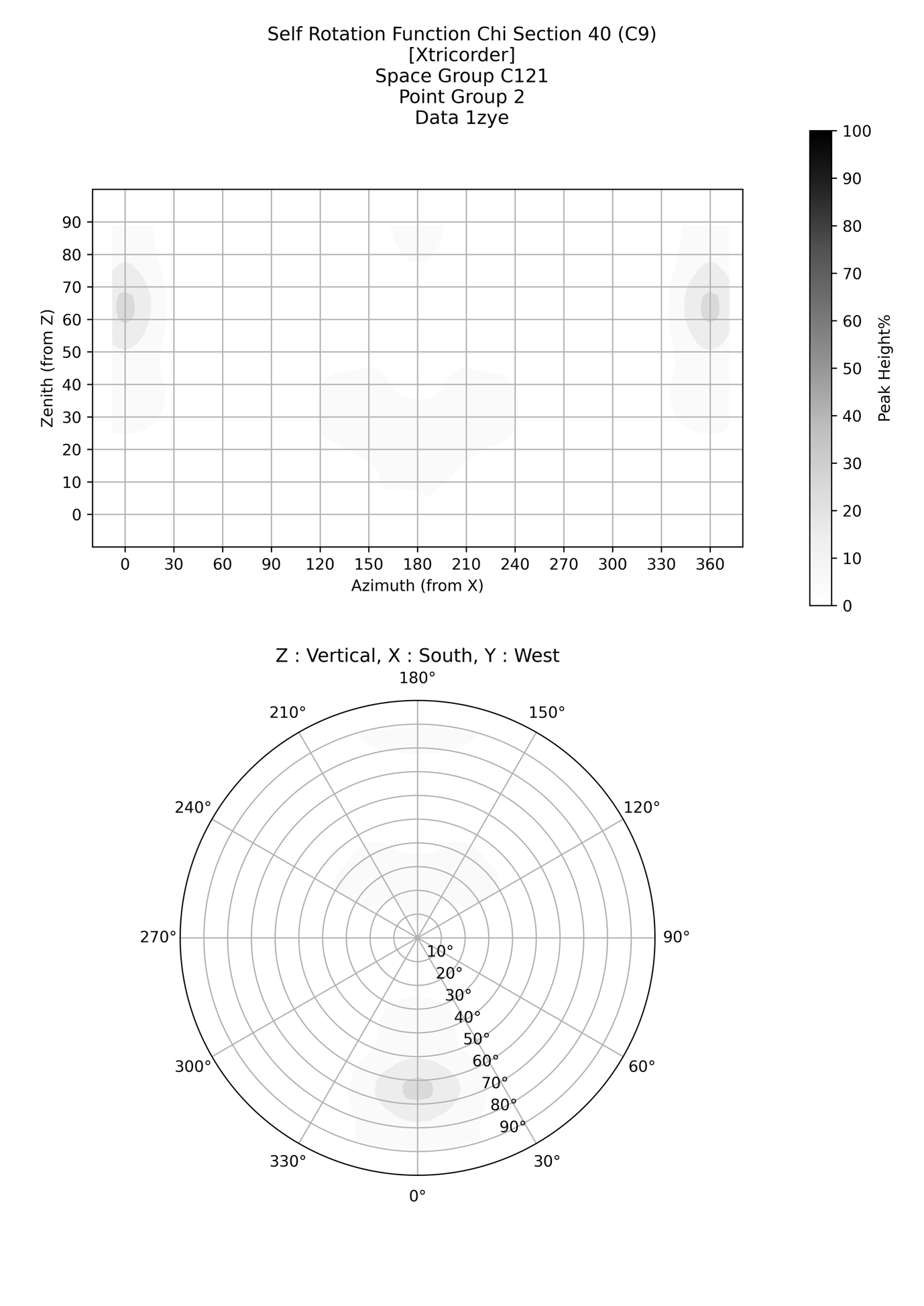

Figure S4 *Stacked SRF images for the protein structure 1ZYE used in machine learning*. A 5x3 grid of greyscale images is shown, each representing a sequential slice from a single orientation stack used by the 3D convolutional neural network during both training and inference. The images encode scatter plots of the SRF peak positions as stereographic projections with marker areas corresponding with peak height. Axes and legends have been removed. Each layer 2-12 contains information about the corresponding rotational section, stars represent crystallographic symmetry and circles non-crystallographic symmetry, with triangles showing that there is a peak near this section but not within the tolerances for be shown as an exact C_n_ rotation. Layer 13 encodes all higher order symmetries as discussed in the text. Layer 1 is a special layer that encodes other relevant parameters and also some SRF information in a different form: in the centre is a space group identifier; Matthews probabilities are encoded at the at the circumference and the presence of circular and dihedral symmetries are encoded on the arc above the point group identifier. Octahedral and icosahedral symmetry is encoded above the point group identifier (if present) and the order of the translational non-crystallographic symmetry is encoded below the point group identifier (if present). Images are shown at the resolution used for training and inference.

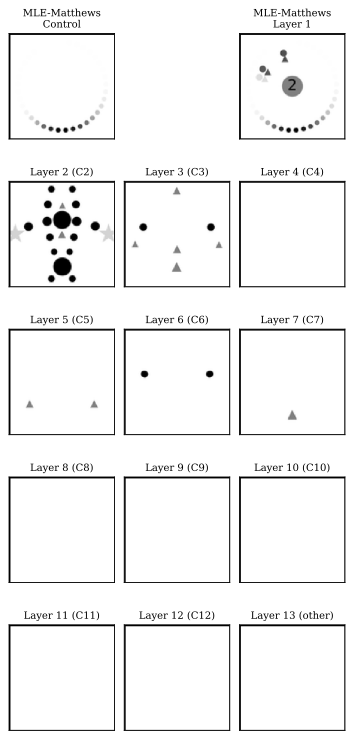

Figure S5 Distribution of structural and crystallographic properties across the dataset. Nine histograms summarizing the frequency of key parameters for structures and their crystallographic data. (a) Crystallographic point group. (b) Solvent content of the crystal lattice. (c) Data anisotropy, represented by ΔB (difference in B-factors along principal axes). (d) Assembly radius, defined as distance from centre of mass containing 90% of atoms. (e) Sphericity of assemblies, defined as the ratio of the longest principal axis to the shortest. (f) Twinning status of the crystallographic data. (g) Number of assemblies per asymmetric unit. (h) Number of residues per chain. (i)  Number of chains per assembly. Each subplot uses logarithmic scaling on the y-axis. Histogram bins and x-axis labels are customized per feature.

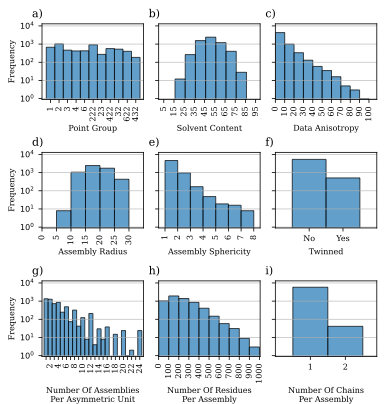

Figure S6 Relationship between symmetry and the number of assemblies per asymmetric unit in dataset. Stacked bar plots showing the distribution of symmetry orders (Cn and Dn) and number of assemblies per asymmetric unit for structures (a, b) For each symmetry order (n in Cn or Dn), the distribution of number of assemblies per asymmetric unit is shown, normalized across all symmetry types. Only n values from 2 to 12 are shown but the proportion of higher order n ≥ 13 values are shown in white. (c, d) For each number of assemblies per asymmetric unit (ranging from 1 to 24), the frequency of associated symmetry orders (n in Cn or Dn) is shown, normalized across all bins. Colours correspond to symmetry order or number of assemblies per asymmetric unit value, as indicated in the legends. Frequencies are normalized within each number of assemblies per asymmetric unit bin (a, b) or symmetry order (c, d). Panel labels (a–d) correspond to each subplot.

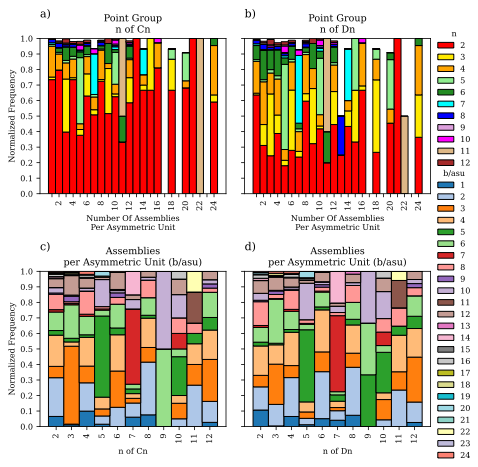
